# Spatial transcriptomic programs relate to spectrolaminar rhythms across macaque cortex

**DOI:** 10.64898/2026.07.08.737187

**Authors:** Ronald Garcia-Reyes, Julien Vezoli, Pedro A. Valdes–Sosa

**Author notes:** These authors contributed equally to this work (Ronald Garcia-Reyes, Pedro A. Valdes–Sosa).

## Abstract

The laminar organization of cortical dynamics is thought to reflect underlying cell-type architecture, but this relationship has not been resolved across primate cortex. Here we developed a spectro-omic framework that allows laminar local field potentials and single-cell spatial transcriptomics to be compared within a common layer-4-referenced coordinate system. Instead of relying on raw band-power analysis, we used Local Spectral Expansion (LSE), a spectrolaminar component model that resolves frequency-by-depth LFP power maps into distinct *δ, θ, α, β*, low-*γ* and high-*γ* components. We then introduced layer-4 projection (L4P) to unfold nonlinear spatial transcriptomic cortical ribbons into flat layer-4-referenced manifolds compatible with the electrophysiological depth profiles. Across twelve matched macaque cortical regions, transcriptomic predictors improved held-out prediction of LSE-derived six-anatomical-layer spectral composition beyond a hierarchy-plus-layer baseline in a partwise logit model: *R*^2^ = 0.621 versus 0.384; Δ*R*^2^ = +0.237; *r* = 0.809 versus 0.692. This gain was supported by hierarchy-preserving shift/reflect nulls 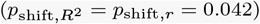 and by a Freedman–Lane region-block residual-permutation test 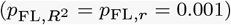. Full-depth PLS1 decoding linked *α*/*β* processes to deep-layer, especially L4/5/6 and L6, glutamatergic programs enriched for axonal, synaptic and myelin-associated biology, whereas low-*γ* and high-*γ* processes were linked to superficial-to-middle-layer GABAergic/PVALB-enriched inhibitory programs together with excitability, ion-homeostasis, non-neuronal and energy-metabolism signatures. Together, these observed spectro-omic relationships suggest that laminar molecular and cellular architecture forms a plausible substrate for the spectrolaminar motif across macaque cortex.

## 1. Introduction

The primate cerebral cortex is a laminated cellular sheet in which local microcircuits, projection patterns and molecular composition vary systematically with depth. This organization has motivated the view that cortical computation is implemented by area-specific variations of a canonical microcircuit^1–6^. It also provides the anatomical substrate for hierarchical communication: supragranular and infragranular neurons differ in their feedforward and feedback projection patterns, and cortical areas vary in cytoarchitecture, projection class and functional specialization^7–12^. Yet it remains unclear whether laminar molecular architecture provides the cellular and biophysical substrate for the depth-resolved electrophysiological rhythms measured at the mesoscopic scale.

This question is now experimentally tractable. Large-scale single-cell and spatial transcriptomic atlases have resolved the molecular and cellular organization of macaque cortex across cortical areas and layers^13^. Complementary resources in marmoset and mouse reveal conserved and species-specialized gradients of cell-type composition, regional identity and gene-expression structure^14,15^. In parallel, laminar electrophysiology has shown that cortical rhythms are strongly organized across depth. Current-source-density analyses have long been used to identify sensory input layers in primate visual cortex^16–18^, and laminar local field potential (LFP) recordings have revealed distinct superficial and deep rhythmic domains in visual, parietal and frontal cortex^19–25^. These findings motivate a direct spectro-omic question: do laminar transcriptomic programs predict the spectral processes expressed across cortical depth?

A key physiological starting point is the macaque spectrolaminar motif described by Mendoza-Halliday et al. Across multiple cortical areas, high-frequency LFP power increases from deep to superficial layers and peaks near layers 2/3, whereas *α*-*β* power increases from superficial to deep layers and peaks near layers 5/6; the *α*-*β*/high-frequency crossover lies close to layer 4^26^. This motif implies that the relevant physiological object is not a single bandpower estimate, nor a set of independent channel spectra, but a joint frequency-by-depth surface, *Y* (*f, z*), in which spectral and laminar structure are coupled.

Most analyses of neural oscillations still begin from predefined frequency bands. This convention has been indispensable for describing superficial high-frequency activity, deep *α*-*β* activity and frequency-specific feedforward– feedback organization. However, fixed-band summaries are limited when spectra contain variable peak frequencies, overlapping rhythms, broadband high-frequency activity, evoked transients, harmonics, non-sinusoidal waveform structure or aperiodic background changes. These concerns were central to the critique by Mackey et al., who argued that broad-band analyses alone cannot establish the presence of ongoing *α*- and *γ*-band oscillations and may conflate narrow-band rhythmic activity with broad-band high-frequency power changes^27^. Thus, even when a spectrolaminar motif is present, the physiological processes that generate it cannot be identified reliably by averaging over fixed frequency windows.

This limitation motivates a process-level model of the full spectrolaminar surface rather than a fixed-band summary. The central issue is whether spectral components should be treated as additive physiological sources or as peaks that modulate a shared aperiodic envelope. Related neural mass, neural field and membrane-biophysics work has shown that cortical oscillations and excitability can be shaped by distributed connectome delays, nonlinear resonance, spatial variation in neural-population density and membrane physicochemical state, reinforcing the need to model spectra as products of spatially embedded dynamical processes rather than isolated band averages^28–31^. *ξ*-*α*NET provides the additive precedent by decomposing spectra into separable latent contributions, whereas FOOOF/specparam-like models fit oscillatory peaks relative to an aperiodic background and are multiplicative in linear power^32,33^. We therefore developed Local Spectral Expansion (LSE) as a laminar additive model in which each frequency-depth map, *Y* (*f, z*), is represented by localized kernels that yield depth-resolved *δ, θ, α, β* and high-frequency profiles for transcriptomic prediction.

A second challenge is laminar alignment. Raw probe contact number is not a cortical-depth coordinate: probes differ in inter-contact spacing, insertion angle, valid-channel count and sulcal orientation. The spectrolaminar motif provides an electrophysiological solution because its *α*-*β*/high-frequency crossover is closely aligned with layer 4. Following the vFLIP-v2 framework, we use this crossover as a layer-4-related anchor and express each probe on a common superficial-to-deep axis, with *z* = 0 at the crossover^34^. This produces a shared laminar co-ordinate for comparing LFP-derived spectral processes across probes, animals and cortical regions. On the transcriptomic side, we apply an analogous layer-referenced transformation, layer-4 projection (L4P), which unfolds spatial transcriptomic cortical manifolds around the empirical layer-4 midpoint and maps cells onto the same depth support as the electrophysiological profiles.

The macaque spatial transcriptomic atlas provides the molecular counterpart to this aligned physiological framework. Chen and colleagues combined large-scale single-nucleus RNA sequencing and spatial transcriptomics across 143 macaque cortical regions, defining 264 transcriptome-resolved cortical cell types and their spatial distributions^13^. This atlas revealed strong laminar and regional structure in glutamatergic, GABAergic and non-neuronal populations, as well as relationships between cell-type distributions and cortical hierarchy. It therefore offers a macaque-specific molecular coordinate system for asking whether regional variation in laminar spectral organization is predicted by the regional and laminar distribution of cell types and gene-expression programs.

Such a comparison requires careful statistical design. Cortical areas vary along anterior–posterior, sensory-association and hierarchical gradients ^7–10^, and transcriptomic cell classes, subclasses and gene-expression programs vary strongly across both area and layer ^13^. Laminar LFP spectra also contain broad areal and depth gradients ^23–26^. A naive correlation between transcriptomic profiles and spectrolaminar power could therefore be inflated by shared geometry rather than by a specific cross-modal relationship. We address this by treating probes and transcriptomic chips as independent samples within matched anatomical regions, estimating regional laminar templates separately for each modality, and testing whether transcriptomic architecture predicts held-out laminar spectral composition. Critically, the primary prediction analysis is evaluated against a hierarchy-plus-layer baseline, allowing us to ask whether transcriptomic laminar architecture explains spectral organization beyond generic cortical-gradient and layer-position effects.

Previous work has linked laminar cell architecture to cortical rhythms, but not at the scale or molecular resolution needed to answer this question. Marker-based studies have related canonical cellular markers to oscillatory power in macaque cortex: parvalbumin (PV), calbindin (CB) and calretinin (CR) are markers of GABAergic interneuron classes and are therefore linked to inhibitory cell programs, whereas neurogranin (NRGN) marks excitatory pyramidal subtypes^35^. This prior work provides an important biological bridge between cellular architecture and oscillatory power, but a key limitation is cellular resolution: a small marker panel cannot resolve the high-dimensional diversity of transcriptomic subclasses, regional cell-type compositions and gene-expression programs captured by spatial transcriptomics. Pathway-level physiological studies have also shown that feedforward and feedback projections can produce cell-type-specific effects on cortical activity^36^. Together, these studies establish important biological priors, but they do not determine whether high-dimensional spatial transcriptomic programs predict the full laminar spectral organization of LFP power across cortical regions.

Here we combine standardized laminar electrophysiology with spatial transcriptomic architecture to test this hypothesis across macaque cortex. We first decompose vFLIP-v2-aligned LFP relative-power maps with LSE to obtain process-resolved laminar spectral profiles. We then project spatial transcriptomic cells into an L4-centered coordinate system and build matched regional laminar templates for cell-type and gene-expression features. Finally, using region-aware transcriptomic decoding with probe/chip bootstrap resampling and laminar null controls, we identify stable cell-type-resolved gene programs associated with distinct LSE-defined spectral processes. This framework tests whether laminar transcriptomic architecture provides a predictive and mechanistically interpretable substrate for regional variation in macaque spectrolaminar organization.

## 2. Results

### 2.1. Local Spectral Expansion defines a laminar spectral atlas

As a first methodological step, we applied Local Spectral Expansion (LSE) to the full macaque laminar LFP relative-power dataset of Mendoza-Halliday et al. and its public Dryad release, rather than only to the subset of regions later used for transcriptomic prediction^26,37^. Each probe-level frequency-by-depth map was vFLIP-v2 aligned to the *α*-*β*/high-frequency crossover, fit with the same six-process additive LSE model, and aggregated into region-level L4-referenced laminar spectral templates.

This analysis converted laminar LFP recordings into a quantitative electrophysiological atlas that could be compared directly with laminar molecular architecture. Raw LFP power is a two-dimensional object, varying jointly with frequency and cortical depth. Because the recordings were aligned by vFLIP-v2 rather than by MRI-guided probe coordinates, all electrophysiological profiles were expressed on a common L4-referenced depth axis, with *z* = 0 at the vFLIP-v2 crossover, negative depths denoting superficial positions and positive depths denoting deep positions.

Local Spectral Expansion (LSE) models each aligned depth-by-frequency power surface as an additive mixture of six canonical spectral processes: *δ, θ, α, β*, low-*γ* and high-*γ* (Fig. 2a). In simplified notation, LSE approximates the observed laminar spectral surface as

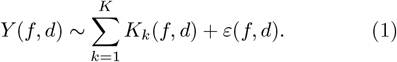

**Figure 1:**
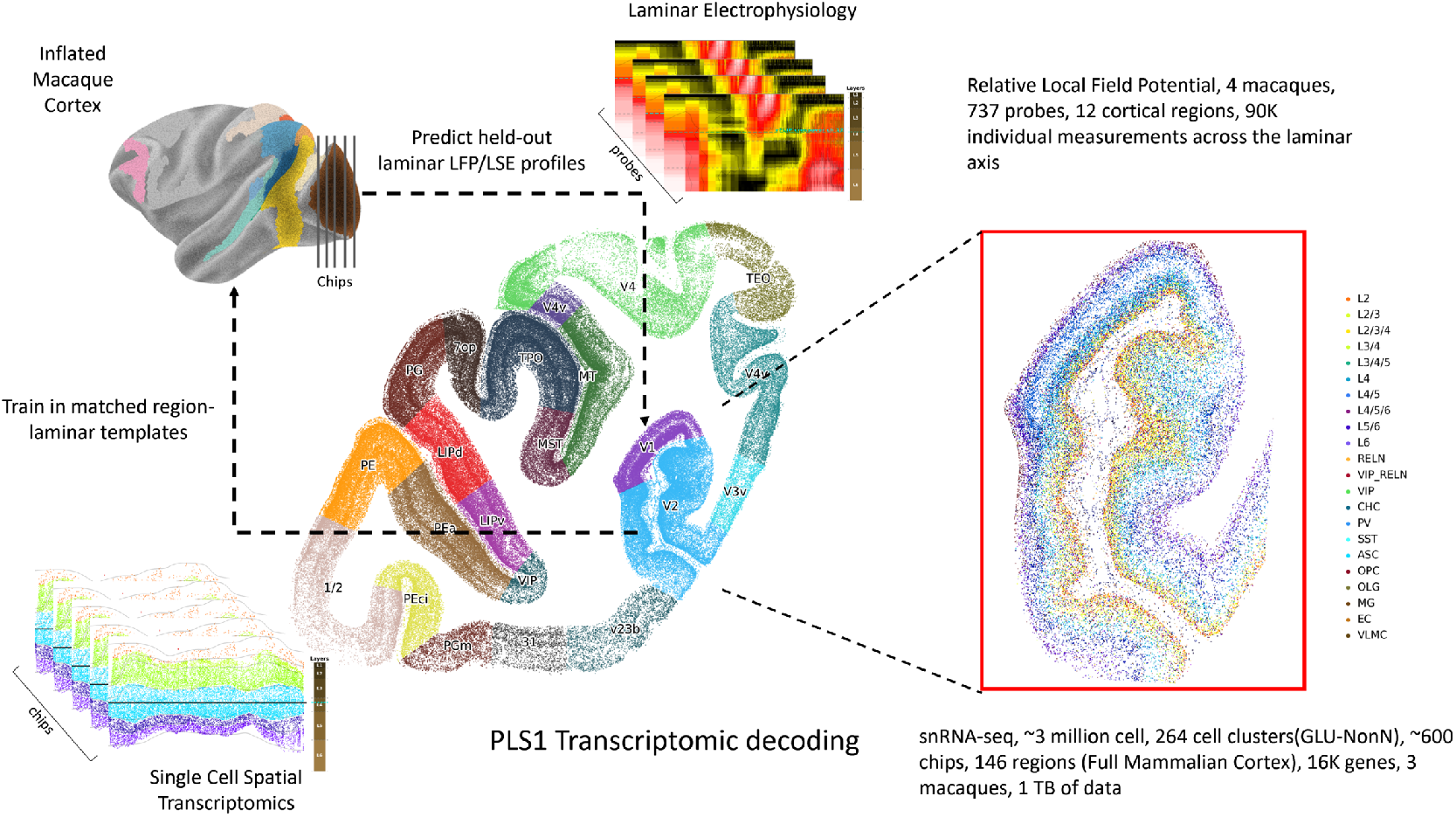
Visual abstract of the spectro-omic prediction framework. **a**, Anatomically matched macaque cortical regions provide the common regional framework for integrating electrophysiological and molecular data. **b**, Single-cell spatial transcriptomic maps capture gene-expression and cell-type profiles across cortical layers within these regions. **c**, Laminar LFP recordings provide frequency-by-depth maps of relative power from the corresponding cortical regions. **d**, Both modalities were aligned on a common layer-4-referenced coordinate system: vFLIP-v2 aligned LFP probes using the *α*-*β*/high-frequency crossover, whereas layer-4 projection (L4P) unfolded transcriptomic cells around the empirical layer-4 midpoint. **e**, LSE converted frequency-by-depth LFP maps into laminar profiles of spectral processes, and spatial transcriptomic gene-expression and cell-type profiles were used to predict these profiles across regions. The primary held-out prediction used a two-component PLS model evaluated by leave-one-region-out cross-validation, whereas full-depth PLS1 decoding was used downstream to identify process-linked transcriptomic programs.

**Figure 2:**
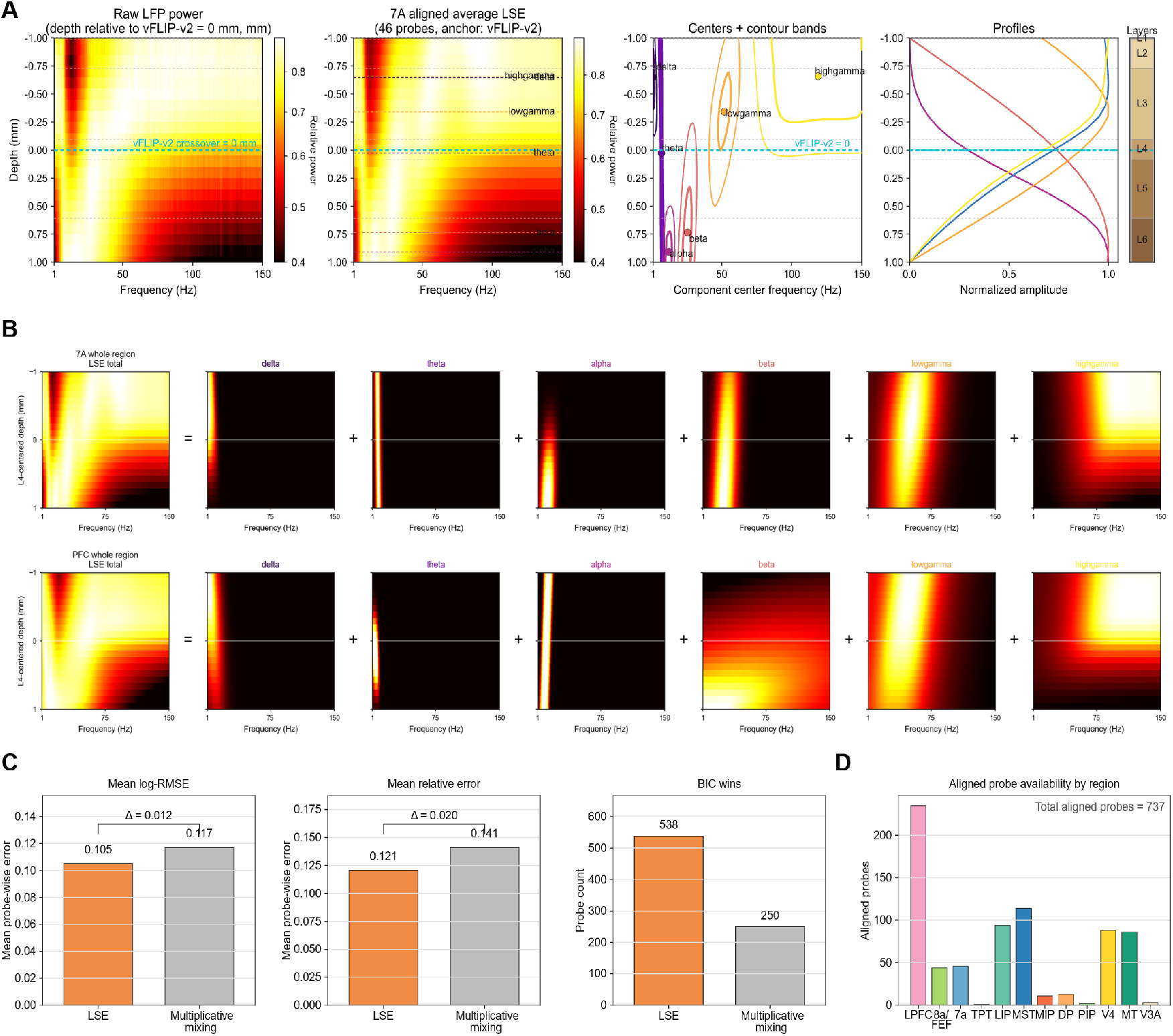
Local Spectral Expansion defines a shared laminar spectral atlas. **a**, Construction of LSE on a vFLIP-v2-referenced laminar depth axis. Left, raw laminar LFP relative power as a function of frequency and depth. Middle, aligned average LSE reconstruction for area 7A. Right, inferred centers, contour bands and normalized depth profiles of the six canonical spectral processes: *δ, θ, α, β*, low-*γ* and high-*γ*. **b**, Additive decomposition of regional spectra into the six LSE processes, illustrating the learned frequency–depth contours for each process in 7A and PFC. **c**, Model comparison between LSE and a multiplicative laminar spectral alternative. LSE showed lower reconstruction error and more frequent BIC support. **d**, Regional availability of aligned probes after vFLIP-v2/LSE quality control. The final electrophysiological atlas contained 737 aligned probes across twelve cortical regions.

Here, *Y* (*f, d*) is the observed relative power LFP surface as a function of frequency *f* and L4-centered depth *d*, and each *K*_*k*_(*f, d*) is a generalized bivariate Lorentzian kernel corresponding to one LSE process. The six kernels correspond to *δ, θ, α, β*, low-*γ* and high-*γ*, and their learned frequency–depth contours can be visualized for each spectral process in Fig. 2b. These Lorentzian kernels are heavy-tailed depth–frequency profiles, allowing each process to have a localized spectral-laminar center while still contributing smoothly across neighboring frequencies and depths. Importantly, LSE does not restrict process geometry to simple ellipsoidal blobs: the learned contours can take more complex shapes, including box-like or plateau-like profiles when supported by the data. This shape information is useful because it models the laminar spread of each spectral process, not only its spectral-laminar localization. The amplitude, frequency center, depth center, spread, orientation and tail behavior of each process are contained within its kernel *K*_*k*_. Details of spectral-component parameter estimation are provided in Methods Section 4.5, and the comparison with the multiplicative spectral model is described in Methods Section 4.6.

This representation preserves the laminar geometry of the signal. Rather than reducing each region to scalar bandpower values, LSE estimates where each spectral process is expressed across cortical depth and frequency. The resulting process profiles form the electrophysiological targets used in the downstream spectro-omic analyses.

In area 7A, whose location on the inflated macaque cortex is shown in Fig. 3a, the aligned LFP spectrum showed the expected macaque spectrolaminar organization, with low-frequency structure concentrated toward deeper compartments and high-frequency structure concentrated toward superficial or middle depths. LSE captured this organization as a sum of interpretable process components, each with its own frequency center, laminar center and spatial spread. Across regions, the same additive framework reconstructed distinct regional spectral surfaces, illustrated for 7A and PFC (Fig. 2b). Thus, LSE provides a shared language for comparing cortical areas: every region is represented by the same six laminar spectral processes.

**Figure 3:**
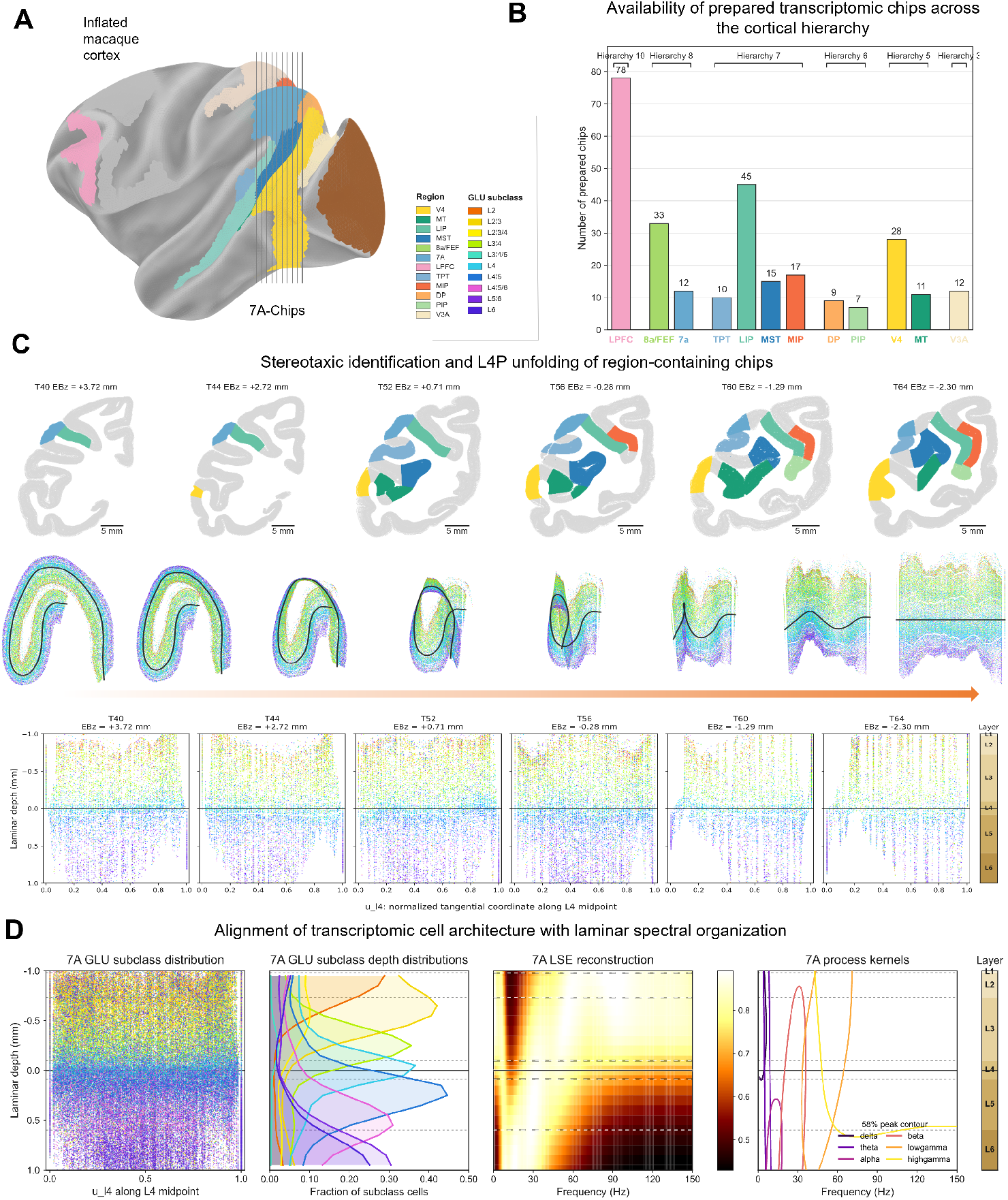
L4P aligns spatial transcriptomic cell architecture with the laminar spectral atlas. **a**, Cortical regions used for transcriptomic chip preparation, shown on an inflated macaque cortex. The 7A chip series illustrates how multiple tissue sections sample one LFP-matched cortical region. **b**, Availability of prepared transcriptomic chips across the cortical hierarchy. Bars indicate prepared region-containing chip annotations before all L4P/QC filtering; downstream predictive analyses used the subset of QC-passing region-chip projections mapped to the common LSE depth support. **c**, Stereotaxic identification and L4P unfolding of region-containing chips. Top, region identities in example 7A-containing sections. Middle, curved cortical ribbons with the empirical L4 midpoint trajectory. Bottom, unfolded L4P coordinates, with tangential position along the L4 midpoint and signed laminar depth orthogonal to L4. **d**, Alignment of transcriptomic cell architecture with laminar spectral organization in 7A. GLU subclass distributions are shown on the L4P manifold, summarized as laminar depth profiles, and compared with the region-level LSE reconstruction and process kernels.

We next tested whether this additive process model provided a better description of laminar spectra than a multiplicative spectral alternative. This model-comparison cohort contained 788 probes for which both LSE and multiplicative fits were available and directly comparable. LSE achieved lower mean log-RMSE and lower mean relative error than the multiplicative model, and was favored by BIC in most probes (Fig. 2c). These results support the use of LSE as the electrophysiological representation for the study: it both improves reconstruction accuracy and yields biologically interpretable laminar process profiles.

After vFLIP-v2/LSE quality control, the final aligned electrophysiological atlas contained 737 probes across twelve cortical regions (Fig. 2d). Probe coverage was uneven across regions, motivating region-level diagnostics that track where sparse or low-support regions are less stable. The LSE atlas therefore provides the electrophysiological half of the spectro-omic bridge: a set of region-level, L4-aligned laminar spectral templates that can be matched to L4P-aligned transcriptomic profiles. Extended Data Fig. 1 shows the broader regional LSE atlas, illustrating how the same LSE procedure was applied across cortical regions to reconstruct frequency-by-depth spectra, decompose them into canonical spectral processes and estimate the laminar profiles used for downstream spectro-omic modeling.

### 2.2. Transcriptomic L4P alignment creates a matched laminar molecular atlas

Having defined LSE-derived laminar spectral targets, we next constructed the matched transcriptomic predictor space from the rhesus macaque (*Macaca mulatta*) single-cell spatial transcriptomic atlas of Chen et al. (Fig. 3)^13^. The Chen atlas provides region- and layer-resolved molecular measurements across macaque cortex, including broad glutamatergic, GABAergic and non-neuronal classes and finer subclasses such as layer-associated GLU groups, PVALB-, SST-, VIP- and LAMP5-related inhibitory groups, and astrocyte, endothelial, microglial, oligodendrocyte, OPC and VLMC non-neuronal groups. These annotations define the molecular variables used down-stream. The electrophysiological and transcriptomic datasets were obtained from different animals, and therefore were matched at the level of species, cortical region and laminar depth, not at the level of individual macaques. Consequently, the analyses below should be interpreted as cross-modal, region- and layer-aligned associations, not as causal evidence linking transcriptomic state to electrophysiological activity.

The key result in this section is geometric. Spatial tran-scriptomic cells are measured in curved two-dimensional cortical sections whose local orientation, cortical thickness and laminar trajectories vary across chips. By contrast, the electrophysiological targets defined above are LSE profiles expressed on a flat L4-referenced laminar axis. Raw Stereo-seq coordinates therefore cannot be compared directly with LSE depth, because such a comparison would conflate cortical curvature, section angle, tissue orientation and true laminar position. The layer-4 projection procedure, L4P, resolves this mismatch by unfolding each transcriptomic cortical ribbon around its empirical layer-4 trajectory, as specified in Methods Section 4.8.

L4P is not simply a visualization transform. It converts each curved transcriptomic ribbon into an L4-centered laminar coordinate system while preserving the empirical laminar organization of cells, subclasses and gene-expression profiles. In this coordinate system, tangential position follows the layer-4 midpoint curve and signed depth is measured orthogonally from that curve. Superficial cells have negative depth, deep cells have positive depth and L4 is centered near zero. L4P therefore plays for transcriptomics the same conceptual role that vFLIP-v2 plays for electrophysiology: vFLIP-v2 places LFP probes on an L4-related flat laminar axis, whereas L4P places spatial transcriptomic cells on the corresponding anatomical L4-centered axis.

Figure 3 summarizes this alignment pipeline. Panel a places the analyzed cortical regions on the inflated macaque cortex and illustrates the chip sampling used for 7A. Panel b shows that prepared transcriptomic chip availability is uneven across the cortical hierarchy, motivating explicit quality control and diagnostic reporting for sparse regions in the all-12 predictive cohort. Panel c shows the core L4P operation: region-containing chips are first identified in stereotaxic tissue sections, then their curved cortical ribbons are unfolded around the empirical L4 midpoint into a common coordinate with tangential position along L4 and signed laminar depth orthogonal to it. Panel d demonstrates why this transformation is useful biologically: after unfolding, GLU subclasses retain the expected superficial, middle and deep laminar organization, and these molecular depth profiles can be displayed on the same L4-centered depth support as the matched 7A LSE reconstruction and process kernels.

The matched transcriptomic subset used here comprised 84 spatial transcriptomic chips from 3 rhesus macaques, mapping 15 omics regions onto the 12 LFP regions. After transcriptomic atlas QC and L4P/LSE laminar projection, the matched atlas contained 4,407,466 projected cells, of which 3,786,877 fell within the LSE-supported depth range. The retained feature table contained 16,183 total features; downstream modeling used 15,912 gene features together with cell-type, subclass and broad-class summaries. Cells were then assigned to the same 21-bin depth grid used for vFLIP-v2-aligned LSE profiles, creating a shared region-depth coordinate for cross-modal analysis.

Thus, the matched object used downstream is not a raw chip overlay and not an artificial one-to-one probe–chip pairing. It is a pair of quality-controlled regional laminar templates: an electrophysiological template derived from probe-level LSE fits and a molecular template derived from L4P-aligned spatial transcriptomics. Region-chip projections were retained only when they contained all six layers, valid midpoint geometry, correct superficial-to-deep orientation, ordered layer medians and L4 centering within the prespecified tolerance. This creates the molecular half of the spectro-omic bridge used in the next analysis, where laminar transcriptomic architecture is tested as a predictor of held-out spectral organization.

### 2.3. Transcriptomic predictors improve laminar spectral reconstruction beyond cortico-laminar gradients

The central prediction analysis asks whether transcriptomic architecture adds information beyond the broad cortico-laminar scaffold introduced in Section 1 and made explicit by the LSE/L4P alignment pipeline in Sections 2.1 and 2.2. Macaque cortical regions differ in hierarchical position, cytoarchitecture, laminar projection patterns and feedforward/feedback organization^7,9,10,36^. Transcriptomic organization also follows large-scale cortical gradients and regional specialization^38,39^. Therefore, a naive transcriptome–spectrum association could be inflated by shared cortical geometry. We tested a stricter question: whether L4P-aligned transcriptomic predictors improve held-out laminar spectral reconstruction after hierarchy and generic layer structure have already been modeled (Fig. 4; Methods Sections 4.10, 4.11 and 4.12).

**Figure 4:**
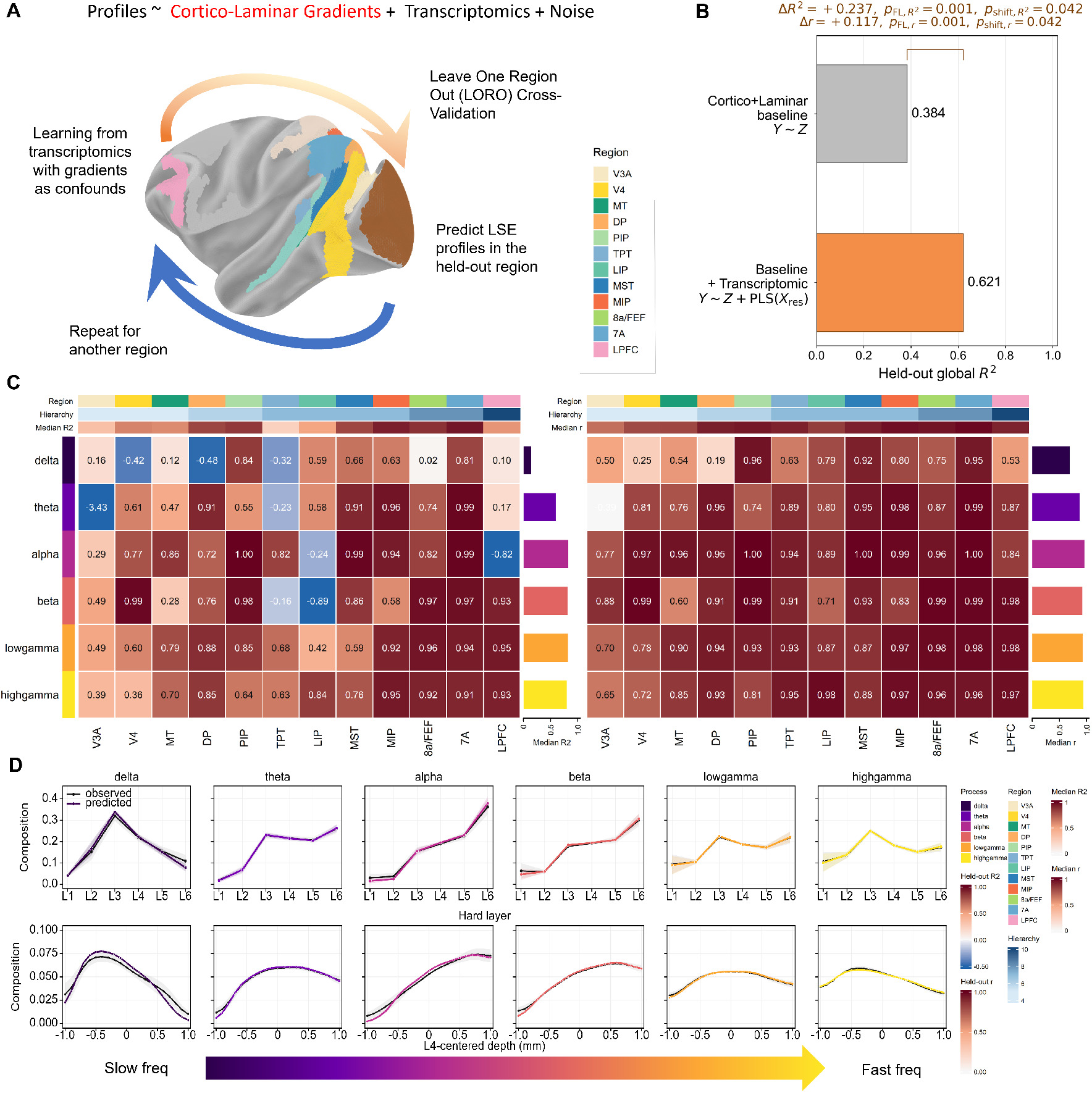
Transcriptomic laminar architecture predicts held-out spectral composition beyond cortico-laminar gradients. **a**, Leave-one-region-out framework. For each fold, eleven regions were used to estimate the cortico-laminar baseline and residualized transcriptomic PLS model, and the twelfth region was held out for testing. **b**, Global held-out performance for the partwise logit hard-layer model after inverse-logit reconstruction and reclosure. The hierarchy-plus-layer baseline reached *R*^2^ = 0.384 and *r* = 0.692; adding transcriptomic information improved performance to *R*^2^ = 0.621 and *r* = 0.809 (Δ*R*^2^ = +0.237, Δ*r* = +0.117). Freedman–Lane region-block residual permutation supported the transcriptomic increment (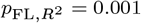, *p*_FL,*r*_ = 0.001), and the hierarchy-preserving shift/reflect null supported exact regional matching (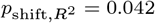, *p*_shift,*r*_ = 0.042). **c**, Region-by-process held-out *R*^2^ and correlation *r* computed on reconstructed layer compositions after inverse logit and reclosure. Individual cells are diagnostics based on six held-out layer values, so negative *R*^2^ can occur even when profile shape remains correlated. **d**, Observed and predicted hard-layer and L4-centered profiles for each canonical process. Predicted profiles recapitulate major laminar motifs in held-out regions, especially for *θ, β* and *γ* processes.

To do this, we performed Leave-One-Region-Out (LORO) cross-validation across all twelve matched regions: V3A, V4, MT, DP, PIP, TPT, LIP, MST, MIP, 8a/FEF, 7A and LPFC. The response was the hard-layer composition of each LSE process, obtained by projecting the shared 21-bin LSE and transcriptomic depth support into six anatomical layers and then applying a partwise logit transform to the layer composition (Methods Section 4.10). The same partwise logit transform was applied to transcriptomic gene-layer compositions, so the reduced and full models were fit in matched transformed layer space. The reduced model contained only cortico-laminar structure, *Y*_logit_ ∼ *Z*, where *Z* denotes the hierarchy coordinate, layer coordinate, hierarchy-by-layer interaction and quadratic terms. We then added transcriptomic information after baseline adjustment, *Y*_logit_ ∼ *Z* +PLS_2_(*X*_res_), where the fixed two-component PLS term was fit to transcriptomic predictors residualized with respect to the same baseline design (Methods Section 4.11). This framing is essential: the goal is not simply to predict laminar spectral profiles, but to test whether transcriptomics explains residual compositional structure after broad cortical gradients and layer geometry are already accounted for.

Figure 4a summarizes this conservative design. In each fold, eleven regions were used to estimate preprocessing, baseline regression and PLS weights, and the twelfth region was held out for testing. The held-out prediction therefore evaluates transfer across cortical areas rather than within-region interpolation. Because the baseline is retained in the full reconstruction and only the residual component is assigned to transcriptomics, improved performance is evidence for spectro-omic coupling beyond the specified confound structure rather than a simple cross-regional regression.

Figure 4b shows the global consequence of adding transcriptomics. The cortico-laminar baseline alone predicted held-out hard-layer composition with global composition-space *R*^2^ = 0.384 and *r* = 0.692, confirming that a substantial fraction of laminar spectral organization is aligned with cortical hierarchy and layer geometry. Adding the transcriptomic term increased held-out performance to *R*^2^ = 0.621 and *r* = 0.809,

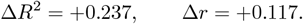

The primary statistical evidence for an incremental transcriptomic effect is the Freedman–Lane region-block residual-permutation test, which supported both the *R*^2^ gain 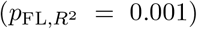 and correlation gain (*p*_FL,*r*_ = 0.001)^40,41^. This test asks whether the transcriptomic PLS term improves prediction after the reduced cortico-laminar model *Y*_logit_ ∼ *Z* has already accounted for hierarchy, layer, hierarchy-by-layer and quadratic structure. In the null, the fitted baseline is preserved, residual responses are permuted by region blocks across the training regions and the transcriptomic term is re-estimated. Thus, Δ*R*^2^ = +0.237 together with 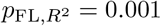 is the main evidence that transcriptomic laminar architecture adds predictive information beyond the specified cortico-laminar baseline.

The hierarchy-preserving shift/reflect null addresses a stricter and different question: whether the transcriptomic gain depends on exact regional matching, or whether a similarly ordered hierarchy-like transcriptomic map would suffice. In the all-twelve-region analysis, 23 unique non-identity shift/reflect mappings were available across the ordered hierarchy, giving a minimum attainable value of 1*/*(23 + 1) = 0.0417. With the empirical formula *p* = (1 + # {*T*_null_ ≥ *T*_obs_})*/*(1 + *N*_shift_), the observed 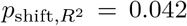 and *p*_shift,*r*_ = 0.042 indicate that the transcriptomic gain exceeded all hierarchy-preserving shift/reflect mappings. This supports the conservative interpretation that transcriptomic structure refines cortico-laminar organization rather than being explained by smooth cortical gradients alone.

Figure 4c shows where this predictive gain is strongest after transforming held-out predictions back from logit coordinates to composition space. The left heatmap reports held-out *R*^2^ for each process–region pair, and the right heatmap reports the corresponding held-out correlation *r*. Prediction was weakest and most heterogeneous for *δ* (*R*^2^ = 0.162, *r* = 0.573), where several region-level cells remained negative. *θ* showed moderate recovery (*R*^2^ = 0.530, *r* = 0.764), whereas *α* (*R*^2^ = 0.777, *r* = 0.893), *β* (*R*^2^ = 0.727, *r* = 0.859), low-*γ* (*R*^2^ = 0.782, *r* = 0.887) and high-*γ* (*R*^2^ = 0.701, *r* = 0.842) were predicted more strongly. The largest transcriptomic increments occurred for low-*γ* and high-*γ*, indicating that faster laminar spectral processes carry the clearest transcriptomic refinement beyond the hierarchy-plus-layer baseline.

Region-level performance was also heterogeneous. Absolute prediction was strongest in 7A, MST, PIP, MIP and 8a/FEF, whereas V3A remained the weakest held-out region despite improvement over its baseline. LIP and LPFC showed sizeable transcriptomic gains beyond base-line even when absolute *R*^2^ was moderate, while DP was the clearest exception in which adding transcriptomics did not improve over the cortico-laminar baseline. These negative or weak *R*^2^ values do not imply absence of laminar structure. Each process–region cell is scored over only six held-out layer values after inverse logit and reclosure, so the full compositional reconstruction can underperform the mean predictor even when observed and predicted profile shapes remain correlated. Taken together, the regional heatmaps indicate that prediction strength is not uniform across cortex: faster *β*, low-*γ* and high-*γ* processes show the clearest transferability, whereas slower *δ* and *θ* organization retains a larger region-specific component.

Figure 4d returns from summary metrics to the predicted profiles themselves. The upper row shows observed versus predicted hard-layer compositions for each process, and the lower row shows the corresponding L4-centered continuous profiles. These reconstructions demonstrate that the model is not merely capturing global process abundance; it recovers the laminar geometry of the profiles. Prediction was particularly tight for *θ, β*, low-*γ* and high-*γ*, where observed and predicted curves closely overlapped. *α* was also recovered in its deepening trend, whereas *δ* showed greater mismatch, consistent with its heterogeneous regional statistics. Thus, transcriptomic predictors recover not only whether a process is present, but how its laminar distribution is shaped across the cortical sheet.

The broader implication is hierarchical. Broad hierarchy and layer geometry establish an important organizational backbone for laminar spectral structure, but they are not sufficient: transcriptomic architecture adds substantial cross-regional predictive information. At the same time, the hierarchy-preserving null shows that this improvement depends on exact regional matching within the ordered cortical map. The most accurate interpretation is that cortico-laminar gradients define the broad scaffold of spectral organization, and transcriptomic programs sharpen, specify and regionally differentiate that scaffold. This transcriptomic refinement is most transferable for *β* and *γ* processes, whereas slow-frequency laminar organization retains more idiosyncratic regional structure.

### 2.4. Functional transcriptomic decoding resolves stable spectro-omic gene programs

#### 2.4.1. Stable PLS1 gene programs

The previous section established the primary predictive result: L4P-aligned spatial transcriptomic architecture improves held-out reconstruction of laminar spectral composition beyond cortico-laminar gradients (Section 2.3). We next asked which molecular programs carry this predictive information. This analysis is intentionally interpretive rather than causal. The electrophysiological and spatial transcriptomic datasets come from different macaques and are matched at the level of species, cortical region and laminar depth (Sections 2.2 and 4.1). Therefore, the full-depth PLS1 analysis identifies stable cross-modal gene programs whose regional-laminar expression covaries with LSE-defined spectral processes; it does not show that those genes generate the rhythms.

For biological interpretation, we used the full-depth 21-bin model rather than the six-layer held-out prediction endpoint. For each spectral process *k*, the response was the residualized region-depth LSE profile and the predictors were standardized region-depth transcriptomic features (Methods Section 4.13). One-component PLS was used because it finds a transcriptomic axis that maximizes covariance with the electrophysiological profile in a high-dimensional, collinear gene-expression space^42,43^. In this setting, PLS1^+^ and PLS1^−^ have directional meanings but not causal meanings. PLS1^+^ genes are the stable genes on the positive side of the transcriptomic axis associated with a spectral process, whereas PLS1^−^ genes are the stable genes on the opposite side of the same axis. With bootstrap loading statistic *Z*_*kg*_ and FDR-adjusted p-value *q*_*kg*_, stable programs were defined as 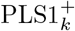 genes satisfying *Z*_*kg*_ > 3 and *q*_*kg*_ *<* 0.05, and 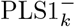 genes satisfying *Z*_*kg*_ *<* −3 and *q*_*kg*_ *<* 0.05. Stability was estimated by resampling regions, LSE probes and transcriptomic chips, aligning the arbitrary PLS signs to the full-data reference and applying FDR control (Methods Section 4.14)^44,45^. Thus, the signed programs should be read as reproducible positive and negative sides of a spectro-omic covariance axis, not as up- and down-regulated genes in the same animal. Figure 5 shows the corresponding process-wise volcano plots for the signed PLS1 gene weights.

**Figure 5:**
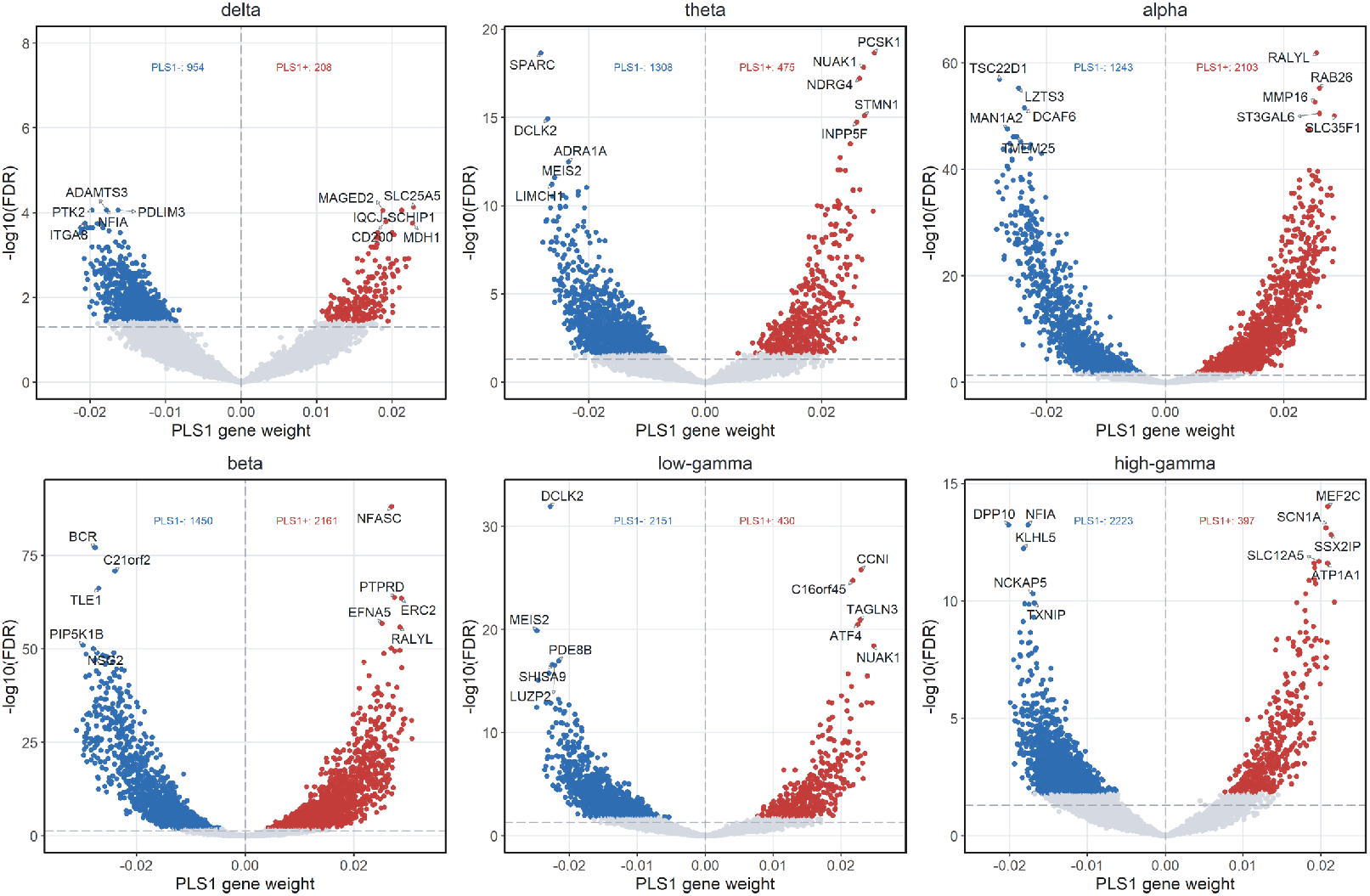
Process-wise PLS1 gene-weight volcano plots. Volcano plots summarize bootstrap-stable gene weights for each LSE process. The x-axis shows PLS1 gene weight and the y-axis shows statistical support, −log_10_(FDR). Red and blue points mark stable PLS1^+^ and PLS1^−^ genes, respectively, whereas gray points indicate genes below the stability threshold. Labels mark representative high-support genes.

The gene-labeled PLS1 axes further separated deep slow-rhythm programs from high-frequency programs (Fig. 5). The *α* and *β* volcano plots showed the clearest convergence between slow LSE processes and structural neuronal programs. The *α* PLS1^+^ side was labeled by *RALYL, RAB26, MMP16, ST3GAL6, SLC35F1, NKD2, SMURF1* and *AFDN*, whereas the *β* PLS1^+^ side was labeled by *NFASC, PTPRD, ERC2, EFNA5, RALYL, AFDN, DLG1* and *MDGA2*. The recurrence of *RALYL* and *AFDN* on the positive side of both *α* and *β* indicates that these deep slow-rhythm axes share part of a common transcriptomic direction. Functionally, this shared axis is consistent with adhesion, membrane organization and synaptic-structural biology, whereas the additional *β* PLS1^+^ labels sharpen the interpretation toward axonal and synaptic architecture: *NFASC* is associated with neurite extension, axonal guidance, synaptogenesis, myelination and neuron–glia adhesion; *PTPRD* is linked to pre- and postsynaptic differentiation; *ERC2* contributes to presynaptic active-zone organization; and *EFNA5* participates in axon fasciculation and guidance^46,47^. Thus, the *α*/*β* programs should be interpreted not only as deep glutamatergic signatures, but more specifically as slow-rhythm covariance axes enriched for adhesion, axonal, synaptic and myelin-associated molecular architecture. The *γ*-linked programs were different. Low-*γ* PLS1^+^ was labeled by *CCNI, C16orf45, TAGLN3, ATF4, NUAK1, FAM241B, SNAP25* and *SATB1*, suggesting a mixed synaptic, cytoskeletal, transcriptional and stress-regulatory program; in particular, *SNAP25* provides a direct synaptic-vesicle and neurotransmitter-release anchor. High-*γ* PLS1^+^ was more directly electrophysiological, with *MEF2C, SCN1A, SSX2IP, SLC12A5, ATP1A1, GABRA1, ATP1B1* and *TENM2* linking high-frequency laminar expression to activity-dependent transcription, sodium-channel excitability, chloride homeostasis, GABAergic signaling and Na^+^/K^+^-ATPase-dependent ion-gradient maintenance^46,47^. The corresponding PLS1^−^ labels, including *C21orf2* on the negative side of both *α* and *β*, and *DCLK2, MEIS2, SHISA9, GSG1L, NFIA, DPP10, NCKAP5, TXNIP, DPYD* and *MSMO1* on the negative side of the *γ* axes, should be read as the opposite sides of stable PLS covariance directions rather than as down-regulated genes or causal suppressors of oscillatory activity. Together, these gene labels refine the cellular enrichment results by suggesting that *α*/*β* rhythms are coupled to structural axonal–synaptic architecture, whereas high-*γ* is coupled more strongly to excitability, inhibitory signaling and energy-dependent ion-homeostasis programs.

#### 2.4.2. Cellular and GO theme summary

Figure 6a gives the functional overview. The alluvial diagram links broad cell classes, enriched subclasses, signed PLS1 programs and dominant GO biological-process themes derived from the enrichment workflow in Methods Section 4.16. Its main message is that LSE-defined activity across the low-frequency to high-*γ* range is associated with both excitatory and inhibitory transcriptomic programs. The *α* and *β* axes are dominated by deep glutamatergic structure, especially L4/5/6 and L6-related GLU subclasses, and connect to synaptic, axonal/projection, myelin and trophic themes. This is consistent with the deep *α*/*β* component of the macaque spectrolaminar motif and with work linking deeper cortical layers to feedback, state maintenance and top-down control^23,26,48–50^. By contrast, *γ*-linked programs pass more strongly through inhibitory subclasses, including PVALB-enriched GABA programs, consistent with the established role of fast-spiking inhibitory circuits in co-ordinating local *γ* rhythms^51–53^. In Fig. 6a, the high-*γ* PLS1^+^ program is also linked to a dominant mitochondrial energy-metabolism GO biological-process theme, indicating that high-frequency spectral structure covaries with cellular energy-demand programs. The alluvial also shows that non-neuronal and extracellular programs are part of the association structure, indicating that LFP spectral organization is coupled to a broader laminar tissue environment rather than to a single rhythm-generating cell type.

**Figure 6:**
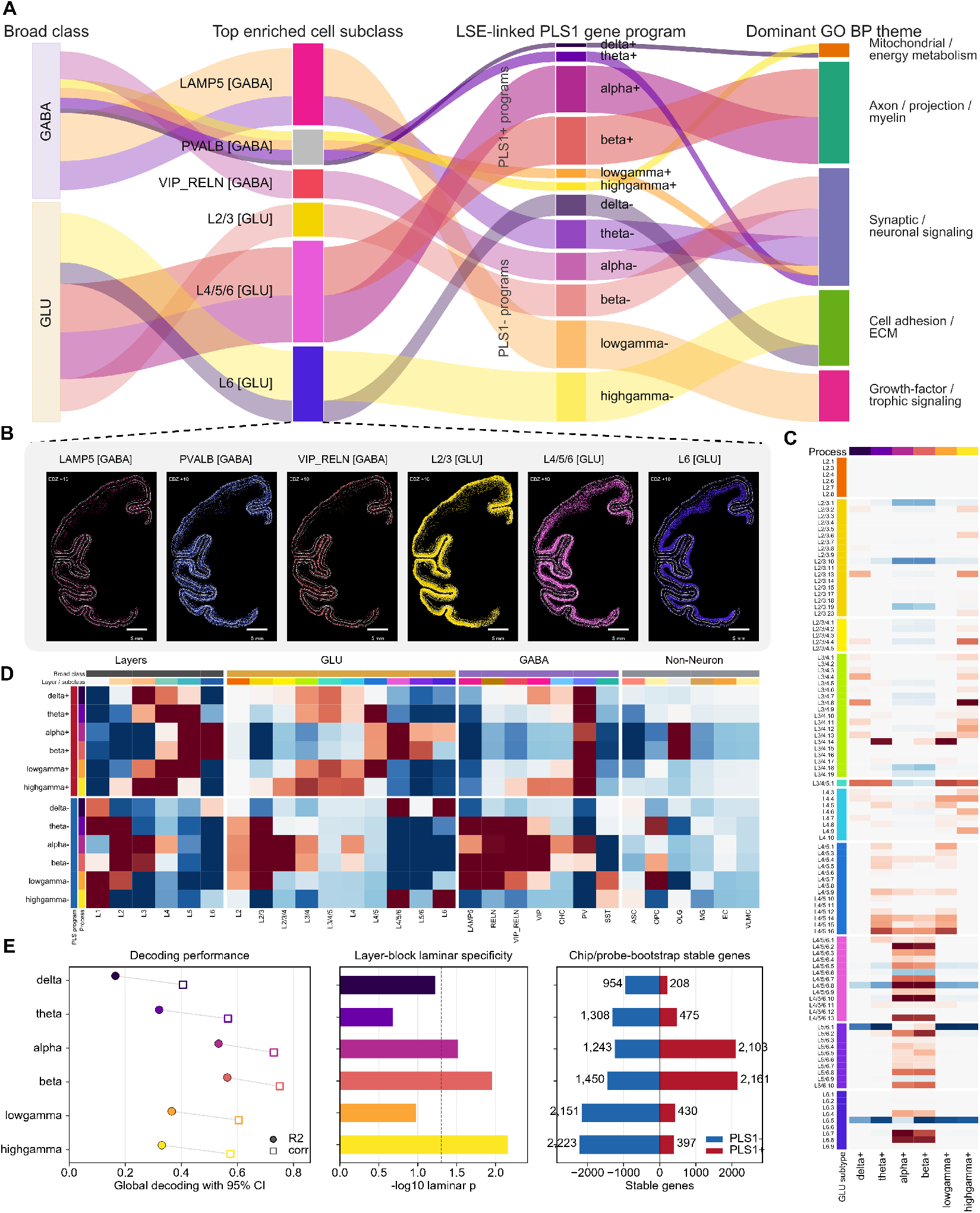
Functional PLS1 transcriptomic decoding of spectro-omic gene programs. Full-depth one-component PLS was used to identify bootstrap-stable positive and negative gene programs associated with each LSE-defined spectral process. **a**, Alluvial summary linking broad cell classes, enriched subclasses, signed PLS1 programs and dominant GO biological-process themes, where theme labels are simplified, manually curated keyword-based summaries of representative enriched GO biological-process terms. **b**, Spatial chip views show representative enriched subclasses on the macaque cortical ribbon. **c**, Fine GLU subclass enrichment for PLS1^+^ programs resolves superficial, middle and deep excitatory contributions across spectral axes. **d**, Integrated signed enrichment across layers, GLU, GABA and non-neuronal annotations separates laminar, excitatory, inhibitory and tissue-environment associations for PLS1^+^ and PLS1^−^ programs. **e**, Full-depth decoding performance, layer-block laminar specificity and bootstrap stable-gene counts summarize model strength and robustness.

Because Fig. 6a is an overview, its dominant GO biological-process labels are deliberately compact. Representative enriched GO biological-process terms were assigned readable keyword-based biological-program names, such as synaptic, projection, trophic, myelin or mitochondrial/energy themes. These alluvial labels therefore provide simplified biological annotations rather than displaying every raw GO term. The complete raw GO biological-process annotations, separated by spectral process and PLS1 sign, are expanded in Fig. 7.

**Figure 7:**
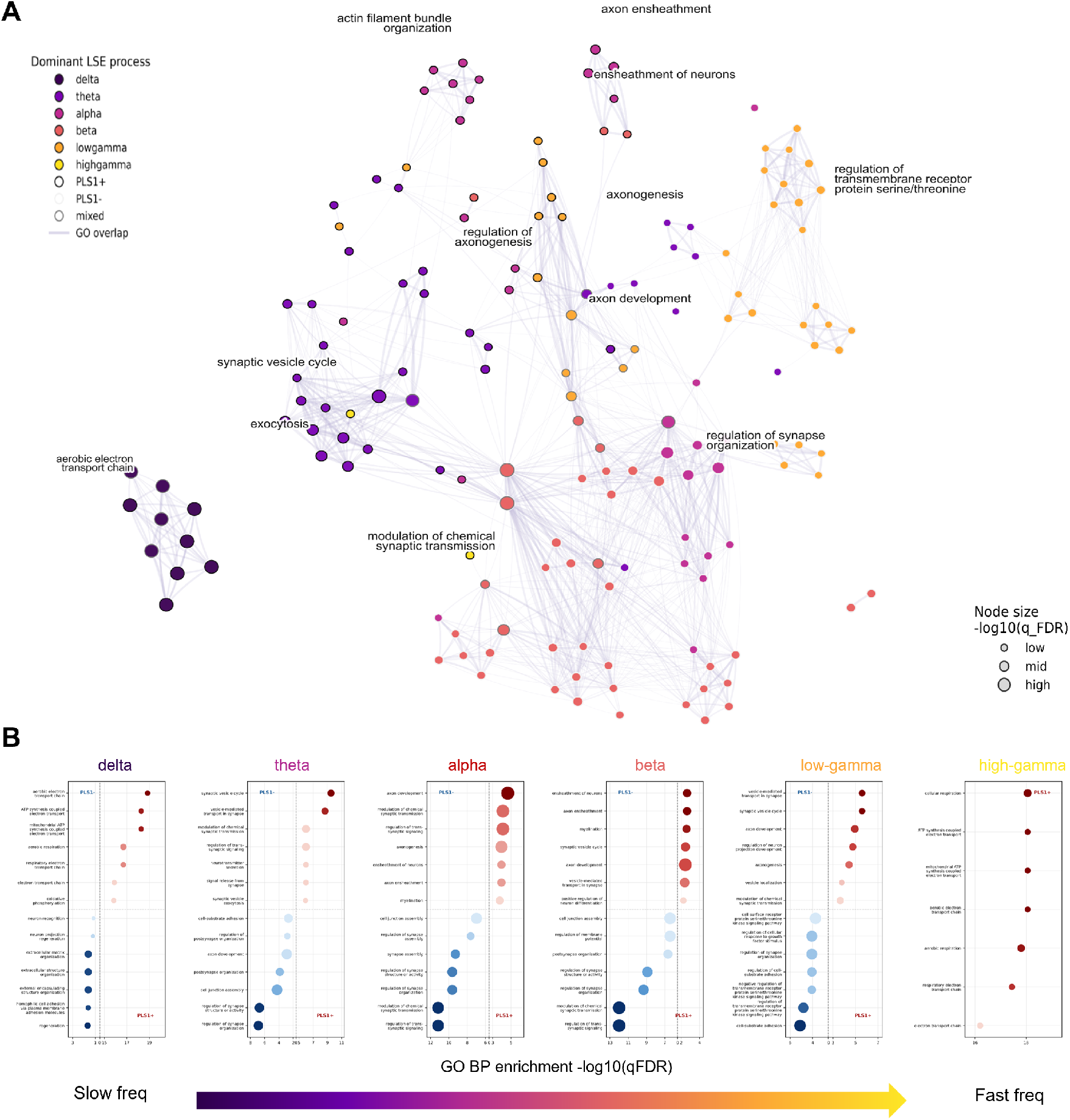
GO BP term-overlap network for stable spectro-omic PLS1 programs. **a**, Functional enrichment network built from stable PLS1 gene programs. Each node is an enriched Gene Ontology biological process (GO BP) term included when it is enriched for at least one stable spectral process/sign gene set, such as *α* PLS1+, *α* PLS1-, *β* PLS1+ or *θ* PLS1-. Node color indicates the dominant LSE process (*δ, θ, α, β*, low-*γ* or high-*γ*), node border indicates the dominant PLS1 sign, and node size scales with GO BP enrichment strength, −log_10_(*q*_FDR_). Edges connect GO BP terms whose contributing stable-gene sets overlap with Jaccard index *J* (*a, b*) = |*G*_*a*_ ∩ *G*_*b*_|*/* |*G*_*a*_ ∪ *G*_*b*_ | ≥ 0.15; the final plotted graph was sparsified by retaining only the strongest few edges per node. **b**, GO BP enrichment was tested using the stable PLS1 gene sets. Raw enrichment *P* values were corrected across tested GO BP terms using the Benjamini–Hochberg false-discovery-rate procedure, and *q*_FDR_ is reported as the adjusted enrichment *P* value.

Figure 6b anchors these enriched subclasses back in spatial anatomy. The displayed LAMP5, PVALB, VIP_RELN, L2/3, L4/5/6 and L6 subclasses occupy distinct positions on the cortical ribbon, illustrating why the L4P transformation was required before transcriptomic programs could be compared with laminar LSE profiles (Section 2.2). These panels are not used as causal evidence; they show that the subclasses highlighted by the enrichment analysis have coherent laminar spatial distributions in the macaque atlas^13^.

Figure 6c resolves the glutamatergic component at higher laminar resolution. Rows are fine GLU subclasses ordered by their laminar position or peak laminar expression, and columns are the PLS1^+^ programs for *δ, θ, α, β*, low-*γ* and high-*γ*. The heatmap reveals a non-monotonic, approximately U-shaped organization along the laminar axis: superficial L2/3–L3/4 subclasses and deep L5/6–L6 subclasses contribute strongly, whereas middle-layer subclasses are more selective. This pattern is important because it shows that the PLS1^+^ axes are not simply generic excitatory-cell signals. Instead, they preserve laminar subclass geometry. The *α*/*β* programs are biased toward deeper GLU subclasses, matching the deep slow-rhythm component of the spectrolaminar motif, whereas *γ* programs include more superficial and mixed laminar GLU structure, consistent with local circuit and feedforward/feedback interactions across layers^20–23^.

Figure 6d integrates the enrichment results across four annotation blocks: anatomical layers, GLU subclasses, GABA subclasses and non-neuronal classes. The layer block confirms that the signed PLS1 axes carry laminar information rather than only regional information. The GLU block separates superficial, middle and deep excitatory programs, with deep GLU enrichment especially prominent for *α*/*β*-associated axes. The GABA block highlights inhibitory subclasses, including PVALB and other interneuron programs, most clearly in the *γ*-linked axes. The non-neuronal block shows that oligodendrocyte, OPC, astrocytic, endothelial or extracellular-environment signals can also be enriched. These signatures should not be interpreted as non-neuronal cells producing oscillations. Rather, they likely index the tissue properties through which LFP fields propagate and are measured, including myelin, axonal architecture, metabolic support, extracellular matrix organization and trophic signaling.

Figure 6e summarizes model strength, laminar specificity and stable-gene counts. Full-depth decoding was strongest for *β* (*R*^2^ = 0.564, *r* = 0.751) and *α* (*R*^2^ = 0.533, *r* = 0.730), whereas *δ* was weakest (*R*^2^ = 0.165, *r* = 0.406). A layer-block null tested whether decoding depended on the correct canonical laminar ordering rather than nonspecific smoothness across depth (Methods Section 4.15). Laminar specificity was strongest for high-*γ* (*p*_laminar_ = 0.0069), followed by *β* (*p*_laminar_ = 0.0111) and *α* (*p*_laminar_ = 0.0306). The stable-gene bars show why the bootstrap criterion is central to the interpretation: *β* had the largest stable program overall (1,450 PLS1^−^ genes and 2,161 PLS1^+^ genes), followed by *α* (1,243 PLS1^−^ and 2,103 PLS1^+^ genes), while low-*γ* and high-*γ* were dominated by stable PLS1^−^ programs (2,151 and 2,223 genes, respectively). These counts identify robust process-linked gene sets for enrichment analysis; they do not imply that every stable gene has an independent mechanistic effect.

Together, Fig. 6 converts the predictive result from Section 2.3 into a biological map. Across LSE components spanning slow activity through high-*γ* up to 150 Hz, stable transcriptomic axes link spectral expression to laminar excitatory, inhibitory and non-neuronal programs. The most parsimonious interpretation is that *α*/*β* activity is coupled to deep glutamatergic and feedback-related architecture, whereas *γ* activity is coupled to inhibitory/PVALB-rich local circuit programs and additional superficial or mixed laminar structure. These are stable, region-depth-aligned spectro-omic associations, not direct causal claims.

#### 2.4.3. Functional enrichment network

Figure 7 provides the detailed functional-enrichment view that complements the compact Fig. 6a alluvial. Whereas Fig. 6a uses simplified, manually curated keyword-based biological theme labels for readability, Fig. 7 displays the raw enriched GO biological-process terms for the stable PLS1^+^ and PLS1^−^ programs of each spectral process. This distinction is important: the alluvial shows the simplified biological theme carried by each process/sign route, while the network shows the underlying ontology terms that support those themes.

In Fig. 7a, each node is an enriched GO biological-process term assigned to the spectral process and PLS1 sign for which it shows the strongest support. Node color identifies the dominant LSE process, the node border identifies whether the term is associated with the positive or negative side of the PLS1 axis, and node size reflects FDR-corrected enrichment strength. Edges indicate that two GO terms share contributing stable genes and were drawn when their contributing-gene sets had Jaccard overlap *J* ≥ 0.15, after retaining only the strongest few edges per node for graph readability. The network should be read as a map of functional redundancy and convergence, not as a pathway diagram: connected clusters imply that several GO annotations are supported by overlapping gene sets, whereas separated modules indicate more process-specific biological annotations. Panel b reports the corresponding FDR-corrected GO enrichment support across process/sign gene sets, making explicit which raw GO terms underlie the broader themes summarized in Fig. 6a.

## 3. Discussion

This study links laminar electrophysiology and single-cell spatial transcriptomics across macaque cortex by placing both modalities on a common layer-4-referenced axis. The principal finding is that laminar transcriptomic architecture predicts held-out LSE-defined spectral composition beyond a hierarchy-plus-layer baseline. The work therefore extends the spectrolaminar motif from a physiological observation to a predictive, cell-type-resolved spectro-omic framework.

The first conceptual contribution is the electrophysiological representation. Mendoza-Halliday et al. established that primate cortex exhibits a reproducible spectrolaminar motif, with superficial high-frequency power, deep *α*/*β* power and an L4-related crossover^26^. We build on that finding by modeling the full frequency-depth surface with LSE. This is necessary because a band average discards the geometry that makes the motif biologically meaningful. LSE assigns each process a frequency center, depth center and shape, enabling an *α* process, a *β* process and *γ* processes to be compared as laminar profiles rather than as broad spectral bins. The additive formulation also matters conceptually: the laminar LFP surface is represented as a superposition of process-like components, making it suitable for comparison with cell-type and gene-program architecture.

The model comparison strengthens this representation choice. LSE outperformed the multiplicative laminar extension of a FOOOF/specparam-like spectral component model in reconstruction error and BIC support, suggesting that, for these macaque laminar LFP maps, the assumption of additive process-like spectral components has stronger empirical support than a model in which oscillatory peaks multiplicatively modulate a shared aperiodic surface. In this sense, LSE extends the one-dimensional *ξ*-*α*NET additive logic into laminar frequency-depth space, while remaining distinct from FOOOF/specparam-style aperiodic-plus-peak models. This does not prove that cortical rhythms are independent circuit generators, but it does indicate that treating *δ, θ, α, β* and *γ*-range processes as separable additive contributors is a useful approximation for this dataset.

The second contribution is geometric. vFLIP-v2 and L4P solve complementary alignment problems. vFLIP-v2 converts probe-native electrophysiology into a flat L4-referenced laminar axis; L4P converts spatial transcriptomic cells from a curved cortical manifold into the same L4-referenced axis. This is not cosmetic. Without vFLIP-v2, the electrophysiological depth coordinate differs across probes. Without L4P, transcriptomic coordinates remain embedded in nonlinear tissue geometry and cannot be directly compared with a flat LSE depth grid. The shared L4 axis is therefore the mathematical object that makes spectro-omic inference possible.

The third contribution is predictive. Transcriptomic laminar structure predicted held-out spectral composition across regions, and prediction remained stronger than a hierarchy-plus-layer baseline. This result is important because cortical hierarchy is both a biological axis and a confound. Macaque cortical areas are organized by laminar projection patterns, and feedforward/feedback pathways are linked to supra- and infragranular compartments^7,9,10,36^. Transcriptomic specialization also follows cortical gradients^38,39^. The hierarchy-controlled model therefore tests a hard question: does transcriptomics add information beyond coarse hierarchy and layer structure? The residual-permutation result supports an incremental contribution, while the hierarchy-preserving null cautions that this contribution is not independent of hierarchy. The safest conclusion is that transcriptomic architecture contributes region-specific predictive information within a hierarchically organized cortex.

The process-specific biology is consistent with, but more resolved than, the existing laminar marker literature. Marker-based work has related canonical cellular markers to macaque oscillatory power: PV, CB and CR index major inhibitory interneuron programs, whereas NRGN marks excitatory pyramidal biology^35^. Our analysis preserves this excitatory–inhibitory logic but moves beyond a small marker panel by resolving layer-specific GLU subclasses, GABA subclasses, non-neuronal classes and signed gene programs.

Within this framework, *α* and *β* were linked most strongly to deep excitatory architecture. The enrichment results point to L4/5/6- and L6-related glutamatergic subclasses, deep L5/6–L6 GLU structure and gene programs enriched for adhesion, axonal/projection, synaptic, myelin-associated and trophic biology. This pattern is consistent with the marker-based findings of Lichtenfeld et al., which linked canonical excitatory and inhibitory markers to macaque oscillatory power, and expands that framework from a small marker panel to spatially resolved GLU subclasses and signed gene programs^35^. It also matches the deep component of the spectrolaminar motif and is compatible with a control-oriented view of *β* rhythms. Task-based frontal-cortex studies provide a cautious physiological context for this interpretation, linking *β* activity to maintained or top-down regimes and *γ* bursts to more transient local activity, but the present analysis does not test task-state stabilization directly^23,48,50^. At the gene-label level, the shared *α*/*β* PLS1^+^ direction included recurrent structural labels such as *RALYL* and *AFDN*, while the *β* axis added axonal and synaptic labels such as *NFASC, PTPRD, ERC2* and *EFNA5*. A deep GLU association therefore suggests that *α*/*β* processes are coupled to infragranular pyramidal architecture, feedback pathways, neuron–glia adhesion, myelin-related structure and integrative dendritic compartments. Johnston et al. further showed in marmoset frontal cortex that deeper *α* activity can suppress upper-layer spiking during proactive saccade control^25^, supporting the idea that deep slow rhythms can regulate superficial activity.

*γ*-linked programs were different and were most naturally interpreted in inhibitory and excitability-related terms. Low-*γ* and high-*γ* processes were linked to superficial-to-middle-layer structure and to GABAergic, PVALB-enriched inhibitory programs, consistent with models and experiments showing that fast-spiking inhibitory networks can generate or pace local *γ* rhythms^51–53^. The low-*γ* PLS1^+^ labels included synaptic and regulatory genes such as *SNAP25*, whereas the high-*γ* PLS1^+^ labels included *MEF2C, SCN1A, SLC12A5, ATP1A1, GABRA1* and *ATP1B1*, linking high-frequency laminar expression to activity-dependent transcription, sodium-channel excitability, chloride homeostasis, GABAergic signaling and Na^+^/K^+^-ATPase-dependent ion-gradient maintenance. This does not mean that *γ* is simply a PVALB marker. The PLS axes also include other inhibitory subclasses, superficial and mixed GLU structure, non-neuronal programs and energy-metabolism signatures. *γ* and high-*γ* in the LFP are mesoscopic signals that reflect coordinated synaptic and transmembrane currents in local tissue, not direct measurements of one cell type. The transcriptomic result should therefore be read as identifying the laminar excitatory–inhibitory and metabolic architecture that predicts *γ* expression across regions.

Non-neuronal programs extend the biological interpretation. Oligodendrocyte-associated signatures argue against an overly narrow rhythm-generator story. Myelin, axonal caliber, extracellular matrix, astrocytic metabolism, oligodendrocyte support and vascular/microglial state all contribute to the cortical microenvironment in which neural fields are generated and propagated. These programs may shape LFP spectra indirectly through conduction delays, synchrony, tissue impedance, metabolic support, axonal density or projection architecture. This interpretation is consistent with previous computational and biophysical work showing that large-scale connectome delays, nonlinear resonance, spatial variation in neuronal density and membrane physicochemical state can shape cortical excitability and oscillatory dynamics^28–31^. In this sense, the paper does not claim that a non-neuronal class generates a rhythm. It claims that laminar spectral organization is predicted by a distributed cellular and molecular architecture that includes neuronal and non-neuronal components.

Several limitations remain. The study is observational and cannot establish causal generation of rhythms by specific genes or cell types. LFP probes and transcriptomic chips are region-matched but not sampled from the same tissue. Several cortical regions have sparse LSE probe support, so their held-out region-level diagnostics are less stable even though they are retained in the all-12 primary cohort. LSE is a structured approximation, not a uniquely identifiable circuit model. The multiplicative comparator provides a useful model benchmark, but additional spectral parameterizations could be tested. The present analysis also treated each probe spectrum primarily as a frequency-by-depth power surface; future work should consider volume conduction within the cortical column and use multivariate probe-level analyses of cross-spectra, coherency or higher-order spectral structure. Higher-order component models such as BiSCA, motivated by nonlinear resonance analyses of cortical oscillations, could test whether nonlinear coupling among LSE-like processes is present in the LFP beyond additive power components^29^. L4P depends on accurate layer annotations and robust estimation of the empirical L4 midpoint. Gene ontology enrichment annotates PLS axes; it does not prove pathway causality. Future work should combine laminar physiology, spatial omics and perturbational approaches in matched tissue, extend the framework across species and tasks, and test whether development, disease or neuromodulatory state alters the spectro-omic mapping.

Overall, these findings support a view in which laminar spectral dynamics are embedded in region-specific cellular and molecular architecture. The cortical column is not only an electrophysiological object and not only a transcriptomic object. It is a joint spectro-omic structure in which deep GLU-associated programs relate to *α*/*β* organization, superficial-to-middle GABA/PVALB-enriched and excitability-related programs relate to *γ* and high-*γ* organization, and non-neuronal tissue programs provide additional context for the mesoscopic fields measured by LFPs.

## 4. Methods

### 4.1. Dataset composition and quality contro

The LFP dataset comprised 942 probes from 4 rhesus macaques (*Macaca mulatta*) across 12 cortical regions: 7A, DP, FEF, LIP, MIP, MST, MT, PFC, PIP, TPT, V3A and V4.^26^ Because these recordings were not acquired with MRI-guided stereotaxic coordinates, analyses were performed using region and laminar depth annotations rather than probe-resolved anatomical coordinates.

Two electrophysiological cohorts were used for different purposes. The additive-versus-multiplicative model-comparison cohort contained 788 probes for which both LSE and multiplicative comparator fits were available and comparable. The final shared laminar atlas cohort contained 737 probes after the later vFLIP-v2 anchor/alignment quality-control steps required to build the common 21-bin L4-referenced atlas. Region-wise passed probe counts in this final atlas cohort were: 7A, 46; DP, 13; FEF, 44; LIP, 94; MIP, 11; MST, 114; MT, 86; PFC, 235; PIP, 2; TPT, 1; V3A, 3; and V4, 88.

Spatial transcriptomic data were derived from the macaque cortex spatial transcriptomic resource of Chen et al., which mapped cell-type organization across rhesus macaque (*Macaca mulatta*) cortex.^13^ The electrophysiological and transcriptomic datasets were obtained from different animals, so they were matched at the level of species, cortical region and laminar depth, not at the level of individual macaques. Therefore, the present study does not support causal inference between transcriptomic organization and electrophysiological activity. Instead, the analyses should be interpreted as identifying cross-modal, region- and layer-aligned associations between laminar transcriptomic composition and laminar LFP spectral organization.

The matched transcriptomic subset used here comprised 84 spatial transcriptomic chips from 3 rhesus macaques, mapping 15 omics regions onto the 12 LFP regions. After transcriptomic atlas quality control and L4P/LSE laminar projection, the matched transcriptomic atlas contained 4,407,466 projected cells, of which 3,786,877 cells fell within the LSE-supported depth range. The retained atlas contained 16,183 total features; downstream modeling used 15,912 gene features together with cell-type, subclass and broad-class summaries.

The full matched LSE-aligned atlas contained 252 region-depth pairs, corresponding to 12 regions sampled over 21 depth bins. The primary predictive analyses used this all-12 matched cohort: V3A, V4, MT, DP, PIP, TPT, LIP, MST, MIP, 8a/FEF, 7A and LPFC. These regions were organized along a discretized macaque cortical hierarchy derived from the visual cortical hierarchy of Felleman and Van Essen and frontal/premotor hierarchy constraints from Cirillo et al.^7,54^. Retaining the full matched atlas allowed the prediction benchmark and the hierarchy-preserving null to test transfer across the complete ordered cortical sequence, while still reporting region-by-process diagnostics that reveal where sparse or low-support regions are less stable. Region names were standardized in the main text as follows: 8a/FEF denotes the FEF mapping to transcriptomic parcels 8AD and 8AV, LPFC denotes the PFC mapping to 46D and 46V, and LIP denotes the union of LIPD and LIPV. The 7A label denotes the electrophysiological 7A region paired to the transcriptomic proxy mapping used in the atlas.

The all-12 predictive cohort corresponded to 252 full-depth region-depth pairs or 72 hard-layer region-layer pairs after reduction to L1–L6.

### 4.2. Spectro-omic analysis framework

We developed a layer-referenced spectro-omic framework to relate macaque laminar local field potential (LFP) spectra to spatial transcriptomic architecture. The analysis combines the laminar coordinate logic of the primate spectrolaminar motif with the region- and layer-resolved structure of the macaque spatial transcriptomic atlas.^13,26^ Electrophysiological probes and spatial transcriptomic chips were treated as independent samples nested within anatomically matched cortical regions. Individual probes were not paired with individual transcriptomic chips. Instead, each modality was first summarized as a regional laminar template on a common layer-4-referenced axis, and cross-modal models were fit between regional electrophysiological and transcriptomic templates.

Let *r* ∈ ℛ index cortical regions, *p* ∈ *p*_*r*_ LFP probes in region *r, c* ∈ *C*_*r*_ transcriptomic chips assigned to region *r, z*_*i*_ aligned cortical depth, *k* an LSE spectral process and *g* a gene or transcriptomic feature. The probe-level electrophysiological observation is *Y*_*prk*_(*z*_*i*_) and the chip-level transcriptomic observation is *G*_*crg*_(*z*_*i*_). Because probes and chips are independent samples, the valid cross-modal object is the pair of regional laminar templates (*µ*_*rk*_(*z*_*i*_), *X*_*rg*_(*z*_*i*_)), rather than an artificial probe-chip pair. Probe-level LSE profiles were modeled as

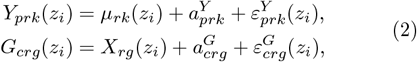

where *µ*_*rk*_(*z*_*i*_) is the regional electrophysiological profile, 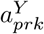 is a probe-level deviation and 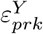 is residual depth-dependent noise. Chip-level transcriptomic profiles were modeled analogously. Regional templates were estimated by robust aggregation within region,

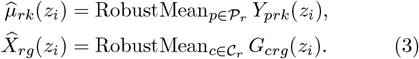

The robust aggregation operator was a pointwise MAD-trimmed mean. At each region-depth coordinate, we computed the median and median absolute deviation (MAD) across probes or chips, retained observations within 3 × 1.4826 × MAD of the median and averaged the retained values. If MAD = 0, the ordinary mean was used; if no value survived trimming, the estimator fell back to the median. All predictive and decoding analyses were performed at the region-depth level, with probe and chip uncertainty propagated by within-region resampling.

### 4.3. Laminar LFP relative-power maps

Laminar electrophysiological inputs were macaque LFP relative-power maps from the public spectrolaminar motif dataset.^26,37^ For probe *p*, the raw spectrolaminar map was denoted *Y*_*p*_(*f, d*), where *f* is frequency and *d* is probe-native depth. The source arrays were organized as probe-by-channel-by-frequency matrices with NaN-padded channel rows. Invalid rows were removed, the remaining channel-by-frequency matrix was transposed for spectral fitting, and the channel coordinate was converted to physical depth using probe-specific inter-contact spacing. Raw channel number was used only as an intermediate coordinate; all regional electrophysiological summaries were constructed after layer-referenced realignment.

### 4.4. vFLIP-v2 electrophysiological alignment

Each LFP probe was aligned to the vFLIP-v2 estimate of the alpha-beta/high-frequency crossover, building on the FLIP/vFLIP family of frequency-based layer-identification procedures.^26,37,55^ For probe *p*, let 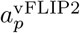 be the probe-specific crossover depth. Probe-native depth was recentered as 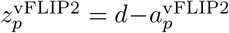, with *z* = 0 at the vFLIP-v2 crossover. Negative values denote superficial positions and positive values denote deep positions. After recentering, each probe was assigned or interpolated onto a common depth support *D* = {*z*_1_, …, *z*_*B*_*}*, where *B*_*z*_ = 21 for the aligned atlas. This common grid enabled cross-probe averaging, region-level LSE template construction and direct comparison with L4-referenced transcriptomic profiles.

### 4.5. Local Spectral Expansion (LSE)

Local Spectral Expansion (LSE) decomposes each laminar relative-power map into localized spectral-depth components. LSE is an additive laminar extension of the *ξ*-*α*NET framework: *ξ*-*α*NET separates one-dimensional spectra into additive latent sources, whereas LSE applies the same additive principle to the two-dimensional frequency-by-depth surface.^32^ The model is additive in linear relative power but optimized in log space, which allows process-specific contributions to be estimated while fitting proportional errors in the observed spectrolaminar map. The formulation was evaluated against a laminar extension of the FOOOF/specparam model, which represents oscillatory structure relative to an aperiodic spectral background.^33^

**Table 1:**
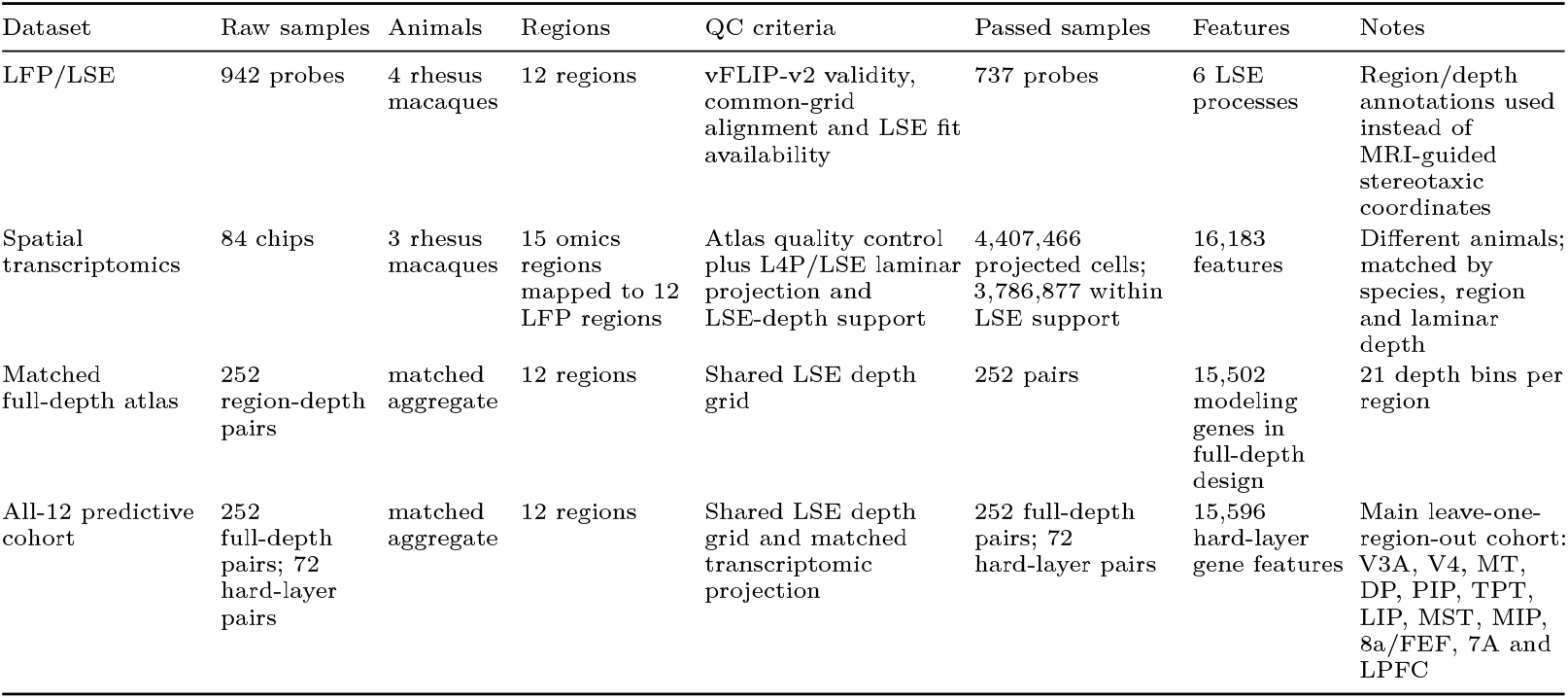
Dataset composition and quality-control summary. Counts are reported for the LFP/LSE electrophysiological resource, the L4P-projected spatial transcriptomic atlas, the full matched region-depth atlas and the all-12 cohort used for leave-one-region-out prediction.

For one aligned probe, LSE approximates the observed surface as 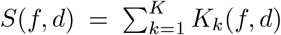, where *K*_*k*_ is a localized component associated with a canonical spectral process. The main process set was *K* = {*δ, θ, α, β*, low*γ*, high*γ}*. The LSE decomposition used six canonical spectral processes: *δ*, 1–4 Hz; *θ*, 4–8 Hz; *α*, 8–13 Hz; *β*, 13–30 Hz; low-*γ*, 30–60 Hz; and high-*γ*, 60– 150 Hz. These supports were used as soft center-frequency penalties rather than hard constraints, allowing each component center to move when supported by the spectrolaminar data.

For component *k*, let *a*_*k*_ > 0 be amplitude, *µ*_*f,k*_ frequency center, *µ*_*d,k*_ depth center, *L*_*k*_ a lower-triangular shape factor and *p*_*k*_ > 0 a generalized-norm exponent. The local displacement, generalized distance and kernel were

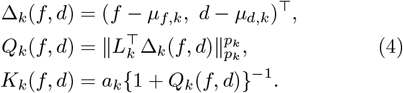

Thus, *L*_*k*_ controls scale, anisotropy and frequency-depth tilt, whereas *p*_*k*_ controls component geometry; *p*_*k*_ = 2 gives the usual elliptical quadratic case. We used

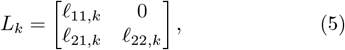

with positive diagonal entries, giving a well-defined local coordinate transform in frequency-depth space.

The production implementation used a mirrored frequency construction inherited from the reference LSE implementation. Each component was represented by positive- and negative-frequency copies with equal half-amplitude: 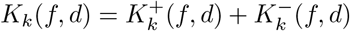 with

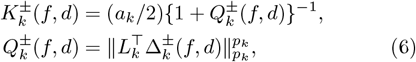

where 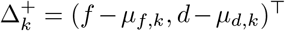 and 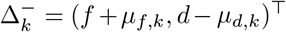. The mirrored copy regularizes low-frequency components while preserving the positive-frequency surface used for fitting.

LSE was fit in log space. For a small stabilizing constant *ε* > 0, *Z*(*f, d*) = log(*Y* (*f, d*) + *ε*) and 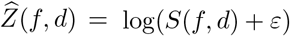. The data term was

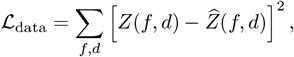

where the sum was taken over finite observations. Component centers were softly constrained to their assigned frequency supports. If component *k* had support bounds 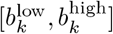, the penalty was *P*_*k*_(*µ*_*f,k*_) = 0 inside the support and increased quadratically outside it. The optimized loss was

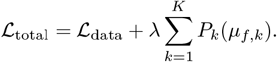

The unconstrained parameter vector for each component was

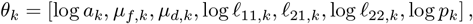

so a *K*-component model had 7*K* unconstrained parameters. For downstream analyses, fitted components were converted into normalized laminar process profiles by frequency marginalization,

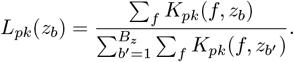

These profiles were the electrophysiological targets used for regional averaging and spectro-omic modeling.

LSE fitting was performed in Python using SciPy’s bounded L-BFGS-B optimizer. Probe-wise spectrolaminar maps were fit independently in log space, with band-initialized LSE components and soft frequency-band penalties. Optimization was stopped at convergence or after 2500 iterations/200,000 function evaluations, and large probe batches were parallelized on a Linux server using 52 CPU workers.

### 4.6. Model scoring and additive-versus-multiplicative comparison

Model residuals were evaluated in log space and model complexity was summarized with the Bayesian information criterion. Let 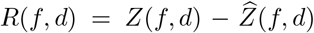 be the number of finite observations and SSE_log_ = ∑ _*f,d*_ *R*(*f, d*)^2^. The maximum-likelihood residual variance estimate was 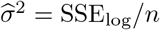. Assuming Gaussian residuals in log space, the log-likelihood was

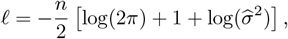

and for parameter count *q*_par_, BIC = *q*_par_ log *n* − 2*ℓ*.

The multiplicative comparator was a laminar extension of the FOOOF/specparam model.^33^ In standard FOOOF, the transformed power spectrum is decomposed into an aperiodic component and Gaussian peak components. We used the same principle on the laminar surface by modeling the transformed relative-power map *Z*_*F*_ (*f, d*) = log*{*1 + *P* (*f, d*)*}* as

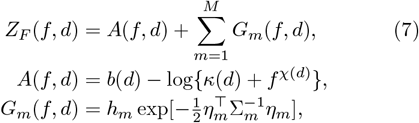

where *A*(*f, d*) is the usual FOOOF aperiodic term, here allowed to vary smoothly with depth through the offset *b*(*d*), knee *κ*(*d*) and exponent *χ*(*d*). The no-knee form is recovered by fixing *κ*(*d*) = 0. The oscillatory term is a multivariate Gaussian peak on the frequency-depth plane, with *η*_*m*_ = (*f* − *µ*_*f,m*_, *d* − *µ*_*d,m*_)^⊤^, amplitude *h*_*m*_ and covariance *Σ*_*m*_. Equivalently, log {1 + *P*_*m*_(*f, d*)} = *G*_*m*_(*f, d*) for each peak, so the peak is Gaussian in the transformed log-power domain and multiplicative after returning to linear power. This defines a physiologically distinct alternative to LSE: LSE treats relative power as a linear superposition of local spectral-depth components, whereas the laminar FOOOF extension treats oscillatory peaks as Gaussian modulations of a depth-dependent aperiodic surface. Both models were evaluated on matched probes using the same log-domain residual and BIC framework.

### 4.7. Spatial transcriptomic chip selection

Spatial transcriptomic chips were assigned to LFP regions using anatomical annotations from the macaque spatial transcriptomic atlas before any electrophysiological comparison.^13^ A chip was included for region *r* if it contained at least one annotated polygon for that region in any cortical layer. Composite LFP-to-omics mappings were defined by anatomical set union; for example, *C*_LIP_ = *C*_LIPD_ ∪ *C*_LIPV_, *C*_FEF_ = *C*_8AD_ ∪ *C*_8AV_ and *C*_PFC_ = *C*_46D_ ∪ *C*_46V_. The 7A electrophysiological label was matched to the transcriptomic proxy mapping used for the corresponding atlas region. This procedure prevented circularity because chip inclusion was determined by anatomical annotation, not by LFP values, LSE parameters, gene-expression similarity or downstream prediction accuracy.

### 4.8. Layer-4 projection of spatial transcriptomic cells

Raw spatial transcriptomic coordinates are not directly comparable across sections because chips differ in curvature, section angle, cortical thickness and visible tissue extent. The layer-4 projection (L4P) procedure was designed for the Stereo-seq macaque atlas geometry, while the layer-resolved pseudobulk logic follows established spatial transcriptomic analyses of cortical laminae.^13,57,58^ L4P expresses each cell by a tangential coordinate along the empirical layer-4 sheet and a signed orthogonal depth relative to the layer-4 midpoint.

For cell *i*, let *x*_*i*_ = (*r*_*x,i*_, *r*_*y,i*_) ∈ ℝ^2^ be its raw chip coordinate. For each chip-region, layer-4 cells were used to estimate a smooth midpoint curve *m*(*s*) ∈ ℝ^2^, parameterized by arclength *s*. The midpoint curve was obtained by rasterizing the layer-4 band, retaining the layer-4-supported connected component, skeletonizing the band, selecting the longest path, smoothing and resampling the centerline, and refining midpoint samples by paired normal ray-casting across the empirical layer-4 mask. Skeletonization followed the digital thinning logic of Zhang and Suen.^59^

For cell *i*, let *m*_*k*_ be the nearest sampled point on the midpoint curve and *S*_*k*_ = ∑_*j<k*_ ∥*m*_*j*+1_ − *m*_*j*_∥ the cumulative arclength to that point. The raw tangential coordinate was *s*_*L*4,*i*_ = *S*_*k*_, and the normalized tangential coordinate was *u*_*L*4,*i*_ = *s*_*L*4,*i*_*/s*_max_, with *s*_max_ = max_*i*_ *s*_*L*4,*i*_. Let *t*_*k*_ be the local unit tangent and *n*_*k*_ = (− *t*_*k,y*_, *t*_*k,x*_) the corresponding normal. The signed raw depth of the cell was 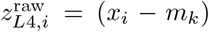. Raw distances were converted to millimeters using the Stereo-seq scale factor *c* = 0.0005 mm*/*unit, so that 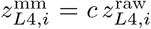. The coordinate convention was negative for superficial positions, zero at the layer-4 midpoint and positive for deep positions.

The normal direction was corrected when necessary using layer medians. If the median raw depth of superficial layers *L*1–*L*3 exceeded the median raw depth of deep layers *L*5–*L*6, the sign of 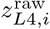 was reversed. A chip-region projection was retained for aligned analysis only when all six canonical layers were present, midpoint construction succeeded, the midpoint was supported by the empirical layer-4 mask, superficial and deep signs were correctly ordered, layer medians followed the expected superficial-to-deep sequence and the layer-4 median depth was close to zero.

### 4.9. Mapping transcriptomic cells to the LSE depth axis

L4P depths were mapped onto the same depth grid used for vFLIP-v2-aligned LSE profiles, linking the histology-validated spectrolaminar anchor to the layer-resolved macaque transcriptomic coordinate system.^13,26,55^ Let 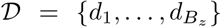 be the electrophysiological depth support. A cell with L4P depth *z*_*i*_ was retained if min(*D*) ≤ *z*_*i*_ ≤ max(*D*) and assigned to the nearest electrophysiological bin *b*(*i*) = arg min_*b*_ |*z*_*i*_ − *d*_*b*_|; its stored aligned depth was 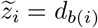. This placed transcriptomic and electrophysiological measurements on an identical L4-centered laminar support.

For chip *c*, depth bin *b* and gene *g*, let *E*_*cbg*_ be the summed expression of gene *g* and *N*_*cb*_ the number of cells in that chip-depth bin. The chip-level mean expression per cell was *X*_*cbg*_ = *E*_*cbg*_*/N*_*cb*_. Regional transcriptomic templates were obtained by aggregating QC-passing chips assigned to each anatomical region.

### 4.10. Projection to hard anatomical layers

For the hard-layer prediction analysis, full-depth electrophysiological and transcriptomic profiles were projected from the 21-bin depth axis to six anatomical layers, analogous to layer-level pseudobulk summaries used in cortical spatial transcriptomics.^13,58^ Let 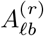 be the region-specific overlap between layer *ℓ* ∈ *{*1, …, 6*}* and depth bin *b*. For an aligned electrophysiological process profile 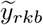, the layer-level signal was 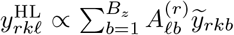 and was then closed to a composition, 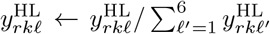. The same region-specific operator was applied to gene-expression profiles to obtain six-layer transcriptomic predictors.

### 4.11. Hard-layer prediction beyond cortico-laminar baselines

The primary confound-controlled prediction analysis used the all-12 matched cohort

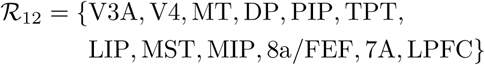

and treated both electrophysiological responses and transcriptomic predictors as laminar compositions, following standard compositional-data principles.^60^ For region *r* and process *k*, the response was a six-part composition *y*_*rk*_ = (*y*_*rk*1_, …, *y*_*rk*6_), with *y*_*rkℓ*_ ≥ 0 and 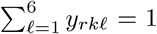. For each transcriptomic gene *g*, the hard-layer predictor was likewise represented as a six-part laminar composition *x*_*rg*_ = (*x*_*rg*1_, …, *x*_*rg*6_). Before model fitting, each compositional component *c*_*ℓ*_ was clipped to avoid boundary values,

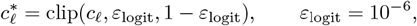

and transformed partwise by

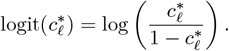

This transform was applied to both response layer compositions and transcriptomic gene-layer compositions. For prediction displays and composition-space metrics, fitted logit values *z*_*ℓ*_ were transformed back by 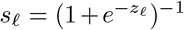 and reclosed,

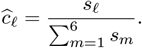

The predictor for region *r* was the concatenated transformed six-layer transcriptomic profile *X*_*r*_ ∈ ℝ^6*G*^.

The hard-layer branch modeled logit-transformed layer values. To remove broad cortico-laminar trends, each region was assigned a fixed discretized hierarchy coordinate using the macaque hierarchy backbone derived from Felleman and Van Essen and Cirillo et al.^7,54^: V3A=3; V4=5 and MT=5; DP=6 and PIP=6; TPT=7, LIP=7, MST=7 and MIP=7; 8a/FEF=8 and 7A=8; and LPFC=10. Each layer was assigned a laminar coordinate *d*_*ℓ*_ ∈ {−2.5, −1.5, −0.5, −0.5, −1.5, −2.5}. Both coordinates were z-scored before model fitting. For each region-layer observation, the baseline design was

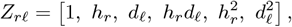

and the cortico-laminar baseline for process *k* was 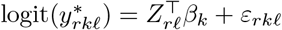.

For the confound-controlled transcriptomic model, both the response and predictors were residualized with respect to the baseline design, *y*_*k*_ = *H*_*y*_*β*_*k*_ + *e*_*y,k*_ and *X* = *H*_*x*_*B*_*x*_ + *E*_*x*_. Equivalently, with the projection *P*_*Z*_ = *Z*(*Z*^⊤^*Z*)^−1^*Z*^⊤^, the residualized objects were *Y*_res_ = *Y* − *P*_*Z*_*Y* and *X*_res_ = *X* − *P*_*Z*_*X*. A fixed two-component multivariate partial least-squares (PLS) model was then fit in residual logit space, 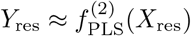, using a latent-variable approach suited to high-dimensional, collinear predictors.^42,43^ For a held-out region, the full prediction was reconstructed as 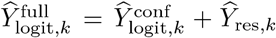. Global performance, region-by-process diagnostics and displayed profiles were computed after inverse logit transformation and reclosure to the simplex.

Prediction was evaluated by leave-one-region-out cross-validation. In each fold, one region was held out, all preprocessing and model fitting steps were estimated from the remaining eleven regions, and the held-out six-layer profile was predicted for each spectral process. The global score pooled 12 × 6 × 6 = 432 reconstructed held-out layer-composition values. Performance was summarized by

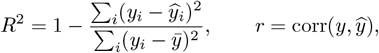

where 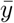 is the mean of the pooled held-out composition values used for the corresponding score. Region-by-process scores were computed over six reconstructed layer values; consequently, individual region-by-process cells can have negative *R*^2^ even when global performance is positive.

### 4.12. Null models for hard-layer prediction

Hard-layer prediction was evaluated with a residual-permutation test for the incremental transcriptomic term and with two region-label null families, following permutation-test logic for empirical significance under structured label reassignment.^40,41^ The incremental contribution of transcriptomics beyond the reduced cortico-laminar model was assessed with a Freedman–Lane residual-permutation null using *N*_FL_ = 1000 permutations. Within each fold and process, the reduced model *Y*_logit_ ∼ *Z* was fit first, residual responses were permuted by region blocks across the training regions, residualized transcriptomic predictors were kept fixed, the PLS model was refit and the held-out region was predicted. This preserves the cortico-laminar scaffold while disrupting the transcriptomic-residual association. The hierarchy-preserving null disrupted exact region identity while preserving coarse cortical ordering by applying non-identity circular shifts and reflected circular shifts to the ordered 12-region sequence. This generated *N*_shift_ = 23 unique mappings. The random region-permutation null reassigned transcriptomic region identities across the twelve regions using 1000 unique non-identity mappings. For a statistic *T*, the empirical p-value was

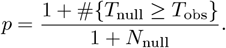

For the shift/reflect family, *N*_null_ = *N*_shift_ = 23, so the smallest attainable empirical p-value was 1*/*24 = 0.0417. This calculation was applied separately to global *R*^2^ and global correlation.

### 4.13. Full-depth PLS1 transcriptomic decoding

The full-depth PLS1 transcriptomic decoding model identified process-specific gene programs associated with LSE laminar profiles on the 21-bin depth axis. PLS was used because it estimates low-rank transcriptomic directions that maximize covariance with the electrophysiological response.^42,43^ This analysis served as the main feature-discovery and biological-interpretation model, whereas the hard-layer model served as the primary held-out confound-controlled prediction analysis.

For each process *k*, the response vector was formed by stacking residualized region-depth LSE profiles, 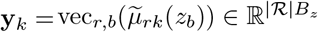. Transcriptomic predictors were stacked into the matrix 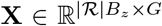, whose columns are standardized gene profiles 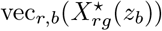. With row-weight matrix Ω = diag(*ω*_*i*_), the one-component PLS direction was chosen as

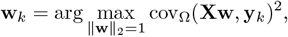

with transcriptomic score **t**_*k*_ = **Xw**_*k*_, scalar regression coefficient 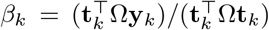 and fitted response 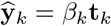.

Weighted correlation and explained variance were computed from the same row weights. For compactness, let 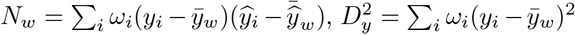 and 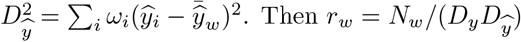 and

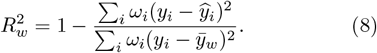

### 4.14. Bootstrap-stable PLS1 gene programs

Stability of full-depth PLS1 gene weights was quantified with *N*_boot_ = 1000 bootstrap replicates, following standard bootstrap inference for estimator stability.^44^ Each replicate resampled regions, LSE probes within sampled regions and omics chips within mapped omics parcels. The aligned electrophysiological and transcriptomic arrays were rebuilt for each replicate, and the one-component residualized PLS model was refit. Because PLS component signs are arbitrary, each bootstrap loading vector was aligned to the full-data reference direction before aggregation: if 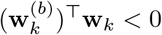, then 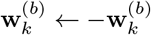.

For gene *g*, the bootstrap standard error was 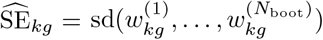. With *w*_*kg*_ denoting the full-data loading, the stability statistic was 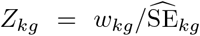. Two-sided G<u>a</u>ussian p-values were computed as 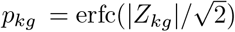 and adjusted within process using the Benjamini-Hochberg false-discovery-rate procedure.^45^ Stable positive and negative process-linked programs were defined as 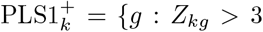 and FDR_*kg*_ *<* 0.05*}* and 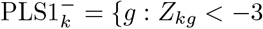 and FDR_*kg*_ *<* 0.05*}*.

### 4.15. Layer-block laminar null for full-depth decoding

Laminar specificity of the full-depth PLS1 model was assessed with a layer-block permutation null, a structured permutation test that preserves within-layer depth-bin order while disrupting canonical laminar alignment.^41^ The 21-bin depth axis was partitioned into six canonical layer blocks, *B* = {*L*1, *L*2, *L*3, *L*4, *L*5, *L*6}. For a null permutation *π* of the six blocks, whole layer blocks were reordered on the depth axis while the sequence of depth bins within each original layer block was left unchanged. The layer-block null used all 6! − 1 = 719 non-identity permutations of the six canonical layer blocks. The model was refit for each null realization, yielding a statistic 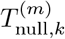, typically 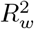. For *M*_null_ = 719 null realizations, the empirical laminar p-value was

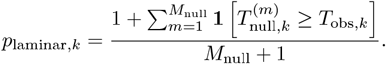

This null tests whether decoding depends on the correct canonical laminar order rather than on region-level expression structure or nonspecific smoothness across depth.

### 4.16. Enrichment analyses

Stable PLS1^+^ and PLS1^−^ gene programs were analyzed against layer, class, subclass, cell-type and pathway gene-set libraries using custom local enrichment code. Cell-type annotations and layer assignments were taken from the macaque spatial transcriptomic atlas, and pathway interpretation used the Gene Ontology (GO) biological-process resource.^13,61^ GO enrichment was run locally with `clusterProfiler::enrichGO` using `OrgDb = org.Hs.eg.db` and `keyType = “SYMBOL”;` therefore, the GO step operated on the harmonized gene-symbol universe passed to `clusterProfiler`. Genes without a valid human-style `SYMBOL` mapping were excluded from GO testing, and the background universe was the saved process-specific gene universe used for the corresponding enrichment run. For each process/sign program, observed overlap with a given annotation set was compared with null overlaps generated by repeated random sampling from the matched gene universe. Empirical p-values were corrected using the Benjamini-Hochberg false-discovery-rate procedure.^45^

Let *S*_*ks*_ denote the stable gene set for process *k* and sign *s*∈ { +,−}, and let *G*_*a*_ denote the gene set for annotation *a*. The observed overlap was *o*_*ksa*_ = |*S*_*ks*_ ∩ *G*_*a*_|. For each random draw *m* = 1, …, *M*, with *M* = 5000, a null set 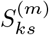 of size |*S*_*ks*_| was sampled without replacement from the same background universe, giving 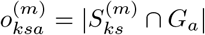. The empirical p-value was

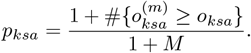

For the corresponding 2 × 2 table, with *a* = *o*_*ksa*_, *b* = |*S*_*ks*_| −*a, c* = |*G*_*a*_| −*a* and *d* = *N* − *a* − *b* − *c*, we computed the Haldane-Anscombe-corrected odds ratio OR_*ksa*_ = (*a*+0.5)(*d*+0.5) */* (*b*+0.5)(*c*+0.5) . Signed enrichment scores were then derived from the odds ratio and the adjusted p-value,

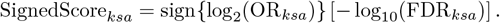

Thus, each enrichment statistic retained both direction and statistical support. These signed scores were used for the GLU subtype heatmap, the global layer plus subclass heatmap and the compact process-by-pathway summaries.

Global cortical-layer enrichment was computed from peak-layer gene assignments using hypergeometric tests with FDR correction. Each gene was assigned to the native layer in which its mean expressio<u>n</u> was maximal, 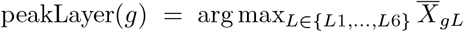. For a process/sign gene set containing *n* genes, if *K*_*L*_ of *N* background genes peaked in layer *L* and *k*_*L*_ selected genes peaked in that layer, the enrichment p-value was *p*_*L*_ = Pr(*X* ≥ *k*_*L*_) for *X* ∼ Hypergeometric(*N, K*_*L*_, *n*). For the Fig. 6a alluvial summary, representative enriched

GO biological-process terms were assigned simplified, manually curated keyword-based biological theme labels; these labels were used only as compact annotations and were not treated as formal semantic-similarity clusters. The exact GO biological-process network in Fig. 7 was built from raw enriched GO terms. Network edges used the Jaccard index between contributing stable-gene sets, *J* (*a, b*) = |*G*_*a*_ ∩ *G*_*b*_|*/* |*G*_*a*_ ∪ *G*_*b*_|, with a threshold of *J* ≥ 0.15 followed by sparsification that retained only the strongest few edges per node. GO biological-process analyses and GO-network visualizations were generated locally using `clusterProfiler, enrichplot` and custom Python/R network scripts, rather than requiring a web-only enrichment platform.^62,63^

## 5. Data and code availability

The laminar LFP relative-power maps and the FLIP/vFLIP-v2 reference implementation are available from Dryad at doi:10.5061/dryad.9w0vt4bnp.^26,37^ The macaque single-cell spatial transcriptomic atlas is available through the Macaque Digital Brain spatial-omics portal at macaque.digital-brain.cn/spatial-omics.^13^ The data used in this research are publicly available datasets; the corresponding ethics approvals and animal-use statements were disclosed in the original macaque LFP and macaque single-cell spatial transcriptomic studies.^13,26^

Custom code for electrophysiological preprocessing, vFLIP-v2 alignment, L4P transcriptomic registration, LSE profile handling, transcriptomic prediction, bootstrap resampling, null-model analyses and figure generation was implemented primarily in `Python 3, R 4.x` and the `MATLAB 2024` release family, with legacy/reference implementations also run in `MATLAB`.^64–72^ The main Python stack used in the current reproducible pipeline includes `NumPy, pandas, SciPy, scikit-learn, matplotlib, DuckDB, NetworkX` and `Pillow`. The main R stack used for heatmaps, enrichment plots and network visualizations includes `ComplexHeatmap, circlize, ggplot2, igraph, ggrepel, patchwork, scales, clusterProfiler, enrichplot, EnhancedVolcano for volcano plots and org.Hs.eg.db`.^62,63,73–76^ The original single-probe LSE and related validation scripts are also available in `MATLAB` form. Scripts used to generate the current analyses are organized in the project repository under `LSE_Spect_vs_Omics`. Exact interpreter, package and toolbox versions, figure-generation scripts, random seeds, input/result roots and archived code-data release metadata will be released with the public repository metadata. Code and metadata are also available from the corresponding authors upon reasonable request.

### 5.1. Computational environment

The computationally intensive analyses were performed on a Linux high-performance server with 104 CPU cores and approximately 257 GB of RAM. The server was used for large-scale parallel jobs that were too expensive for the local workstation, including LSE model fitting across probes and regions, transcriptomic atlas construction, chip-level spatial projection, pseudobulk aggregation, and bootstrap/permutation analyses.

## 7. Acknowledgements

We thank Professor Eduardo Martinez-Montes for useful discussions. We also thank Professor Mu-Ming Poo for useful insights that helped shape this research.

## Appendix Extended Data

**Extended Data Fig. 1.**
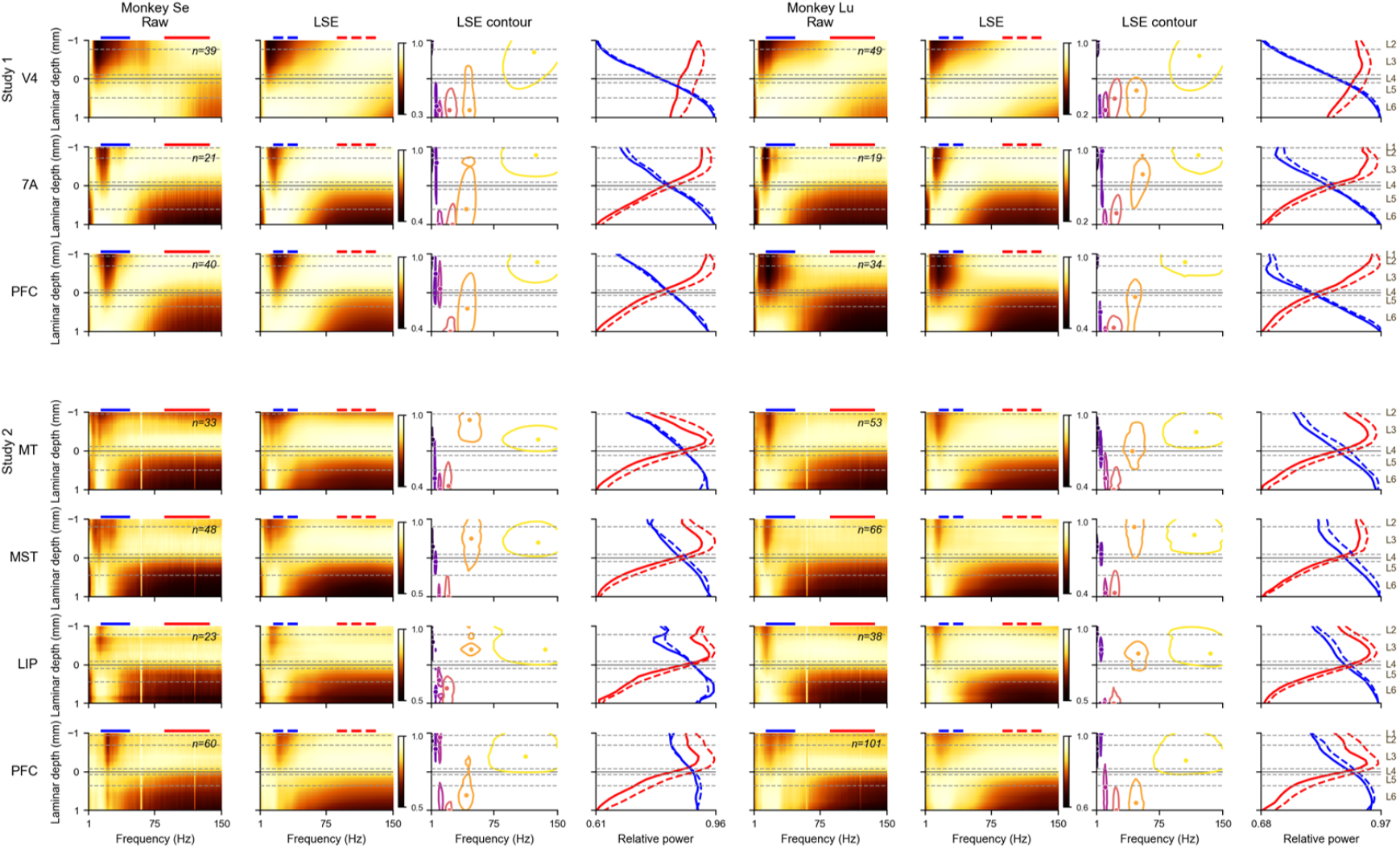
Regional LSE atlas grid. Regional aligned LSE templates and component profiles across the broader electrophysiological atlas. For each selected cortical region, probe-wise LSE fits were averaged after vFLIP-v2 alignment to form the displayed region-level LFP/LSE spectrum and its decomposition into *δ, θ, α, β*, low-*γ* and high-*γ* processes. The plotted depth coordinate is the normalized L4-referenced frame from *z* = −1 to *z* = 1, with *z* = 0 at the L4/vFLIP-v2 anchor, negative values superficial and positive values deep. The *α*/*β* and high-*γ* profiles generally cross near this anchor, indicating agreement between LSE-estimated spectral organization and the vFLIP-v2 alignment reference. Sparse regions are retained in the all-12 leave-one-region-out prediction cohort, and their lower support is reflected in the region-level diagnostics rather than handled by exclusion.

